# The utility of environmental data from traditional varieties for climate-adaptive maize breeding

**DOI:** 10.1101/2024.09.19.613351

**Authors:** Forrest Li, Daniel J. Gates, Edward S. Buckler, Matthew B. Hufford, Garrett M. Janzen, Rubén Rellán-Álvarez, Fausto Rodríguez-Zapata, J. Alberto Romero Navarro, Ruairidh J. H. Sawers, Samantha J. Snodgrass, Kai Sonder, Martha C. Willcox, Sarah J. Hearne, Jeffrey Ross-Ibarra, Daniel E. Runcie

**Affiliations:** Department of Evolution and Ecology, University of California Davis, Davis, CA 95616, USA; Department of Plant Sciences, University of California Davis, Davis, CA 95616, USA; Center for Population Biology, University of California Davis, Davis, CA 95616, USA; Institute for Genomic Diversity, Cornell University, Ithaca, NY 14853, USA; United States Department of Agriculture-Agricultural Research Service, Robert W. Holley Center for Agriculture and Health, Ithaca, NY 14853, USA; Department of Ecology, Evolution, and Organismal Biology, Iowa State University, Ames, IA 50011, USA; Department of Molecular and Structural Biochemistry, North Carolina State University, Raleigh, NC 27607, USA; Laboratorio Nacional de Genómica para la Biodiversidad/Unidad de Genómica Avanzada, Cinvestav, Irapuato, México and Program in Genetics, North Carolina State University, Raleigh, NC 27695-7614, USA; Department of Plant Science, The Pennsylvania State University, State College, PA, USA; International Maize and Wheat Improvement Center (CIMMYT), Carretera Mexico-Veracruz, KM45 Carretera Mexico-Veracruz, El Batan, Texcoco, Estado de Mexico, 56237, Mexico; Genome Center, University of California Davis, Davis, CA 95616, USA

**Keywords:** local adaptation, environment-of-origin, gene-environment association, genomic prediction, traditional variety diversity

## Abstract

Maintaining crop yields in the face of climate change is a major challenge facing plant breeding today. Considerable genetic variation exists in *ex-situ* collections of traditional crop varieties, but identifying adaptive loci and testing their agronomic performance in large populations in field trials is costly.

Here, we study the utility of climate and genomic data for identifying promising traditional varieties to incorporate into maize breeding programs. To do so, we use phenotypic data from more than 4,000 traditional maize varieties grown in 13 trial environments. First, we used genotype data to predict environmental characteristics of germplasm collections to identify varieties that may be locally adapted to target environments. Second, we used environmental GWAS (envGWAS) to identify genetic loci associated with historical divergence along climatic gradients, such as the putative heat shock protein *hsftf9* and the large-scale adaptive inversion *Inv4m*.

Finally, we compared the value of environmental data and envGWAS-prioritized loci to genomic data for prioritizing traditional varieties. We find that maize yield traits are best predicted by genomic data, and that envGWAS-identified variants provide little direct predictive information over patterns of population structure. We also find that adding environment-of-origin variables does not improve yield component prediction over kinship or population structure alone, but could be a useful selection proxy in the absence of sequencing data. While our results suggest little utility of environmental data for selecting traditional varieties to incorporate in breeding programs, environmental GWAS is nonetheless a potentially powerful approach to identify individual novel loci for maize improvement, especially when coupled with high density genotyping.

## Introduction

Protecting crop and wild plant populations against the harmful effects of climate change is one of the most important goals of plant genetic research today. Global temperatures have increased 1.5°C above pre-industrial levels (Hulme 2016), with further increases of 3.3-5.7 °C possible in the next 100 years (Intergovernmental Panel On Climate Change (IPCC)2023). Agricultural systems have already been impacted by warming (Lesk *et al*. 2016), and models under future warming scenarios indicate this trend will be exacerbated, with yield losses across major staple commodities in the range of 25-50% by 2100 (Challinor *et al*. 2014).

One promising strategy for maintaining crop productivity under climate change is to harness existing genetic variation to identify sources of abiotic stress resistance (Varshney *et al*. 2018;Hellin *et al*. 2014; Dwivedi *et al*. 2016). Traditional domesticated varieties (*i*.*e*., landraces) of many crops have been evolving in diverse environments for thousands of years (Bellon *et al*. 2018; Brush 1995; McCouch 2004), resulting in a broad base of genetic variation and adaptation to a wide range of environmental niches. Farmer seed networks and *ex situ* conservation efforts over the past century have preserved extensive collections of traditional varieties not just for preserving genetic diversity in crops, but also for desired trait qualities (McLean-Rodríguez *et al*. 2021; Hellin *et al*. 2010; Ramirez-Villegas *et al*. 2022).

Incorporating diversity from traditional varieties into breeding programs for quantitative traits such as yield is a difficult endeavor. Traditionally, breeders have first identified traditional varieties with apparently useful characteristics, then backcrossed these against elite germplasm to introgress adaptive alleles while maintaining the agronomic performance afforded by the genetics of elite material in the rest of the genome (Bernardo and Yu 2007; Mayer *et al*. 2020; Snowdon *et al*. 2021). Because this is difficult to do comprehensively with tens of thousands of traditional varieties, the Focused Identification of Germplasm Strategy (FIGS) was proposed to prioritize traditional varieties based on their environment of origin (Mackay 1990; Khazaei *et al*. 2013). In FIGS, a sample of traditional varieties are first grown in field trials to identify beneficial traits, then trait presence is modeled as a function of environmental variables to identify other traditional varieties in the full collections that may possess similar traits. FIGS has been successfully applied to identify drought adaptation in broad bean, flower frost tolerance in wild strawberry, and seed traits in bread wheat (Khazaei *et al*. 2013; Egan *et al*. 2018; Kehel *et al*. 2020).

A limitation of FIGS is that it does not consider genetic information. If beneficial exotic alleles are not specifically tracked and prioritized using genetic markers, they are rapidly lost when backcrossed into elite material (Yang *et al*. 2020; Allier *et al*. 2020). Early use of molecular markers in breeding used QTL mapping and marker assisted selection to select candidate varieties and introgress specific alleles (Stuber *et al*. 1982; Tanksley *et al*. 1989). With the increased accessibility of whole genome marker data, genomic selection approaches have become widely used to make selections among candidate varieties (Jannink *et al*. 2010; Bernardo and Yu 2007). Now that large numbers of accessions in crop germplasm repositories are being sequenced, there is the opportunity to use genomic prediction to prioritize accessions for use in breeding programs (Crossa *et al*. 2016; Gorjanc *et al*. 2016). For instance, Yang *et al*. (2020) proposed Origin Specific Genomic Selection (OSGS) as a way to maintain potentially beneficial exotic alleles during line development. Implementing genomic selection using diverse traditional varieties introduces many challenges, including collecting sufficient high quality training data on important traits in relevant environments – thus, it has yet to be known whether this approach will be more successful than FIGS.

As the environment-of-origin data underlying FIGS and the genome-wide marker data underlying genomic selection may be complementary, a selection approach that uses both data could provide additional advantages (Lasky *et al*. 2015; Kehel *et al*. 2020). Environmental and genomic marker data can be combined to identify specific loci with allele frequencies that differ along environmental gradients. If populations are locally adapted to their environment, then the frequencies of loci underlying environmental adaptation may show correlations with environmental gradients. Genome-wide scans for alleles with frequencies correlated to environmental gradients are called environmental genome-wide association analysis (envGWAS) (Lasky *et al*. 2023), and have been applied in natural populations such as Loblolly pine and *Medicago* as well as traditional varieties of crop species such as maize, barley, and soybean (Frichot *et al*. 2013; Yoder *et al*. 2014; Romero Navarro *et al*. 2017; Anderson *et al*. 2016; Bandillo *et al*. 2017). The combination of environment and genomic data should be especially important for evaluating varieties in future environments. Lasky *et al*. (2015) showed that Sorghum alleles associated with drought and low pH successfully predicted variation in plasticity in field trials.

Maize (*Zea mays spp. mays*) is the most productive crop globally and has adapted to grow in nearly every environment that humans inhabit (Hake and Ross-Ibarra 2015), yet yield in modern maize lines has already been impacted by climate change (Rosenzweig *et al*. 2014). Domesticated maize varieties and wild teosintes are well adapted to their distinct local environments (Lafitte *et al*. 1997; Bellon *et al*. 2018; Mercer *et al*. 2008; Fustier *et al*. 2019; Pyhäjärvi *et al*. 2013) and constitute an extensive but largely untapped well of genetic diversity for plant breeding (Liu *et al*. 2016; Prasanna 2012). For example, traditional maize varieties grown in common gardens across elevation gradients in Mexico demonstrated substantially higher fitness when grown in trials more similar to their native environments than when grown in non-local trials (Mercer and Perales 2019), and macrohair and anthocyanin traits showed evidence of adaptive divergence across highland/lowland environment in reciprocal transplant experiments using a broader population of North and South American accessions (Janzen *et al*. 2022). Local adaptation in maize and teosinte has resulted in selection on a number of individual loci (Pyhäjärvi *et al*. 2013; Fustier *et al*. 2019), many of which show strong associations between genotype and the abiotic environment, including inversions such as *Inv4m* associated with flowering time (Romero Navarro *et al*. 2017; Gates *et al*. 2019) and *Inv9f* associated with macrohair growth (Calfee *et al*. 2021), as well as individual genes such as HPC1 that regulate phospholid metabolism and flowering time (Barnes *et al*. 2022).

Maize may therefore be an ideal candidate for a strategy that combines modern detection methods of adaptive variation with novel technologies to minimize the challenges of using exotic germplasm. CIMMYT (the International Maize and Wheat Improvement Center) curates the world’s largest collection of over 24,000 traditional maize varieties, a resource that could play a crucial breeding role in mitigating the negative effects of climate change on maize. With such a large and diverse collection, narrowing down favorable material adaptive to target climates to introduce into pre-breeding programs is a difficult task.

In this study, we identify adaptive genetic variation in a core set of CIMMYT’s collection of maize germplasm. We evaluate methods for jointly using genomic and environmental data to estimate the utility of environmentally associated alleles in predicting the genetic value of traditional varieties. First, we show that traditional maize varieties are locally adapted along multiple environmental gradients using large-scale field trials and by measuring the genome-wide correlation between genetic variation and environmental gradients. Next, we perform envGWAS to identify individual genetic markers associated with environmental gradients and find dozens of potentially adaptive loci. Finally, we evaluate the utility of envGWAS-identified markers and environmental data for prioritizing useful traditional varieties compared to conventional genomic prediction methods. We find that envGWAS candidate loci explain little additional variation for yield component traits compared to population structure and genome-wide kinship, but are enriched for phenotypic associations in field trial data, providing opportunities for prioritized introgression via methods such as OSGS. We also find that environment-of-origin data alone can provide modest predictive ability, which may be useful for large germplasm collections that have not yet been comprehensively genotyped or phenotyped.

## Results

### Phenotypic trials demonstrate local adaptation in maize

To be able to test different approaches for identifying genetic diversity useful for agronomic improvement, we used a panel of *≈* 4, 100 traditional maize varieties from CIMMYT’s maize seed bank (Figure 1). Of these, GBS genotyping (Elshire *et al*. 2011) and growing-season environmental data from their collection location were available for 3511 genotypes representing 2895 seed bank accessions, which were used for genotype-environment associations. We combined these data with phenotypes from the CIMMYT Seeds of Discovery (SeeD), consisting of a set of 23 common garden trials representing 13 locations over multiple years at varying elevations (Gates *et al*. 2019; Gorjanc *et al*. 2016; Romero Navarro *et al*. 2017, see Supp. Table S1). We evaluate phenotypic data from crosses with common testers in multiple common gardens across Mexico, totaling more than 280,000 progeny and spanning almost 18,000 plots (see **Materials and Methods**). We focused on four yield components (see Table S3: grain weight per hectare (GWPH), field weight (FW), plant height (PH), and bare cob weight (BCW), measured on 144 to 1434 (mean = 586) testcross families per trial (Supp. Table S2). In total, testcross families representing 2518 accessions that had genotype, climate, and phenotype data were used for phenotypic analysis.

**Figure 1.**
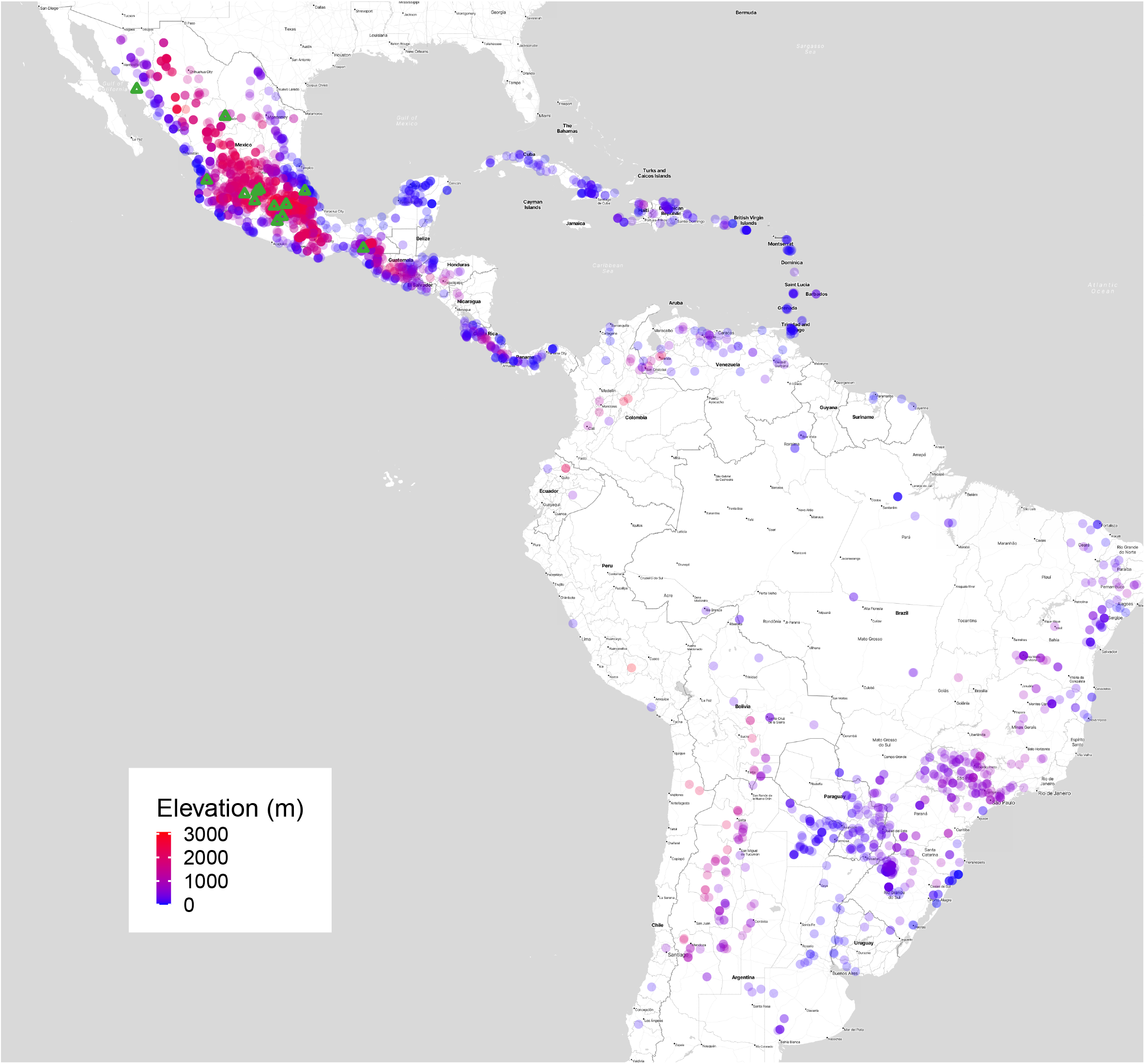
Map showing the 2895 unique geo-referenced locations of traditional maize variety accessions from CIMMYT used for envGWAS, colored by elevation. Green triangles in Mexico indicate locations where field trials were held.

To test for local adaptation in our sample of traditional varieties, we modeled the correlations between yield-related traits of a traditional variety and the annual environmental values of its collection location (Figure S4). If the traditional varieties are locally adapted to an environmental gradient, we predicted that the environment-of-origin values corresponding to the highest yield should be positively correlated with trial environmental values. We fit quadratic curves to yield-related trait values along three environmental gradients (elevation, mean annual temperature, and total annual precipitation) and estimated the slope of the regression of the apex of these curves on the environmental value of each trial.

For all yield traits, the posterior mean of the slope parameter was positive for elevation, precipitation, and temperature, indicating that adaptation predictably occurs along these environmental clines (Supp. Figure S5). 95% posterior intervals of the slope parameter included the value *β*_*h*_ = 1 for field weight along elevation ([0.34, 1.22]), precipitation ([0.17, 4.07]), and temperature gradients ([0.02, 1.13]), indicating that for field weight our results are consistent with a one-to-one relationship between trial environment and estimated optimal environment, providing strong evidence of local adaptation along all three clines (Figure 2). For grain weight per hectare, posterior means of the slope were smaller than one, but still positive, providing moderate evidence for local adaptation along these environmental clines. However, posterior interval widths for this trait were much larger, because this trait was measured in fewer trials.

For bare cob weight and plant height, while we observe positive posterior means for slope for all environmental variables (Supp. Figure S5g-l), 95% posterior intervals were not consistent with a slope of one, save for plant height along precipitation ([0.20587697, 1.9036396]), indicating that while there is a correlation between the optimal environment and the true environment for these traits, traditional varieties may be under-adapted for these traits to the range of environments we tested here. Interestingly, we see that the posterior mean slope for plant height on precipitation is close to one (Supp. Figure S5k), but the intercept was much higher than zero. This suggests that the optimal precipitation for a trial for maximizing plant height is always much higher than the actual trial’s precipitation value.

**Figure 2.**
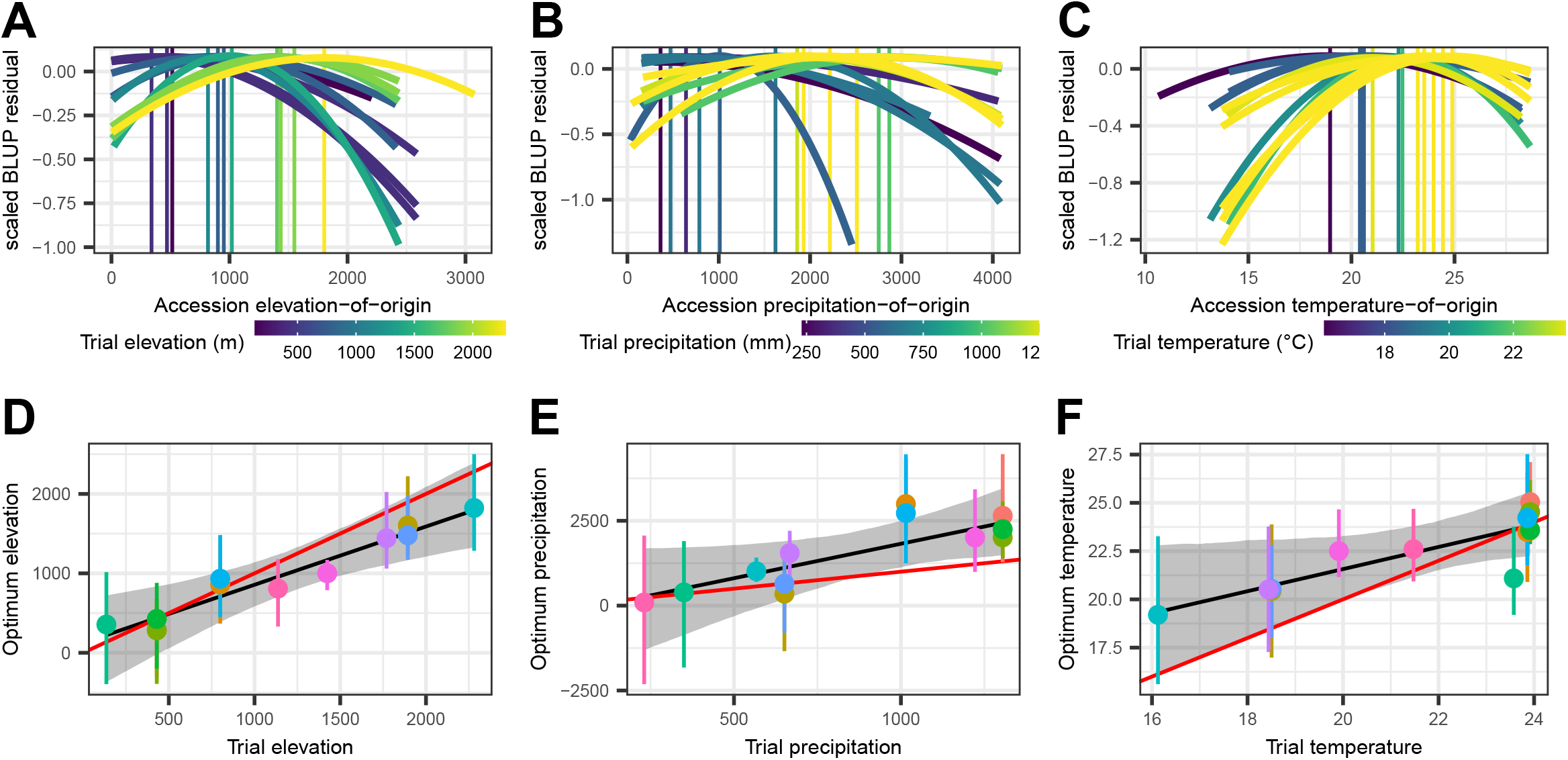
Phenotypic trials show evidence of local adaptation for field weight along elevational and climatic clines. **A-C)** Transfer distance plots showing phenotype values for each maize accession as a function of environment-of-origin. Measures of elevation **(A)**, annual total precipitation **(B)**, and mean temperature **(C)** are represented on x-axis. Each quadratic curve represents a posterior model for scaled field weight BLUP residuals of all accession testcross families grown in a given trial against environment-of-origin value, and colored by that trial’s environmental value. Vertical lines represent the optimal environment value for that trial as determined by the model. **D-F)** Posterior probabilities of the optimal environment value for a given trial against real value for the trial environment. Points represent individual trials, with whiskers representing the 95% interval of posteriors for that trial’s optimum. The black line and grey ribbon show posterior mean and 95% posterior intervals of the relationship between optimal environmental value and the trial’s environmental value. The red line shows a 1-1 linear relationship for comparison.

### Genotypic variation across environment provides further evidence of local adaptation

If traditional varieties are locally adapted as suggested by our trial data, we should likewise observe broad scale correlations between genotype and environment. To test this, we created a dataset of growing season environmental variables for 2895 unique collection locations (see **Materials and Methods**). We then matched these environmental data to samples with available genotyping by sequencing (GBS) data for a total of 3511 genotypes (Romero Navarro *et al*. 2017) at 345,270 imputed biallelic SNPs with minor allele frequency *>* 0.01.

To directly estimate how well genotype data alone predicts environmental variation among accessions, we built a genomic prediction of environment (GPoE) model for all environment variables using MegaLMM (Runcie *et al*. 2021) and evaluated its accuracy using 10-fold cross-validation (Figure 3). GPoE predictive ability was high for elevation (average Pearson’s *r* = 0.839), temperature range (*r* = 0.838), and minimum temperature (*r* = 0.829), and moderate for other variables (0.637 < *r* < 0.761; see Figure 3).

**Figure 3.**
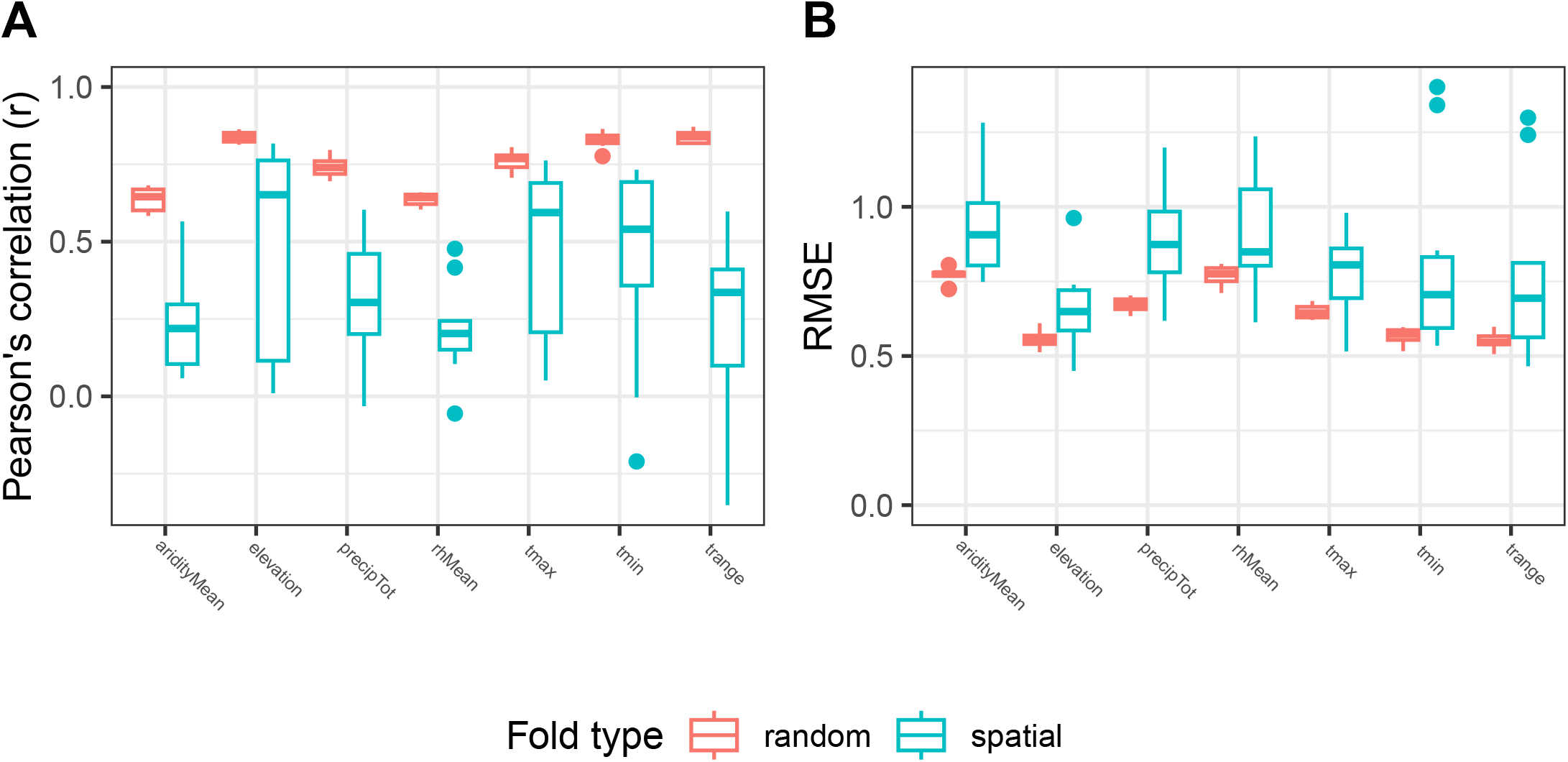
Correlations between genetics and environment are demonstrated through prediction but decrease when considering spatial distance. For each environmental variable, the Pearson’s *r* correlation **(A)** and root mean square error (RMSE) **(B)** between observed and predicted environment values of random (salmon) and spatial (cyan) CV GPoE models.

While local adaptation predicts a strong correlation between genotype and environment, an alternative explanation is that environmental variation is spatially autocorrelated at a scale greater than typical dispersal. Under such a scenario, even in the absence of adaptation, individuals would be found in environments highly similar to their relatives, and thus environments of collection would be predictable from genetic relationships. To distinguish these possibilities, we split our study population into 10 spatially separated sub-samples (see **Materials and Methods**) and repeated the above GPoE analysis, predicting environmental values of individuals from one geographic region using only individuals from other geographic regions to train our models.

Predictive ability for spatially distant accessions is still greater than zero for all environmental variables (*P* < 0.001) but is significantly lower (*P* < 0.001) and has higher RMSE (*P* < 0.001) and higher variation among folds than for randomly sampled accessions (Figure 3).

### Gene-environment association identifies candidate loci

Despite the lower predictive ability of spatially explicit models, our GPoE analysis points to a meaningful association between genetic variation and environmental variation. To identify loci contributing to this relationship, we employed a multitrait environmental GWAS (envGWAS) implemented in JointGWAS (Runcie 2022) to find SNP markers associated environmental variation.

Our envGWAS model controls for residual correlation between environmental factors and population structure and prioritizes SNPs associated with at least one of the environmental variables (see **Materials and Methods**).

We found 330 SNPs with a p-value less than 10^*−*5^ (Figure 4A), which we grouped into 32 independent loci based on LD. For most of these candidate loci, linkage disequilibrium decays within several Kb, with notable exceptions of *Inv4m*, a large 13Mb structural variant located on chromosome 4; the *hsftf9* locus at SNP *S*9_148365695, with significantly associated SNPs 250kB away in LD; and SNPs at *S*9_101922947 and *S*10_41085126 (Supp. Figure S11).

**Figure 4.**
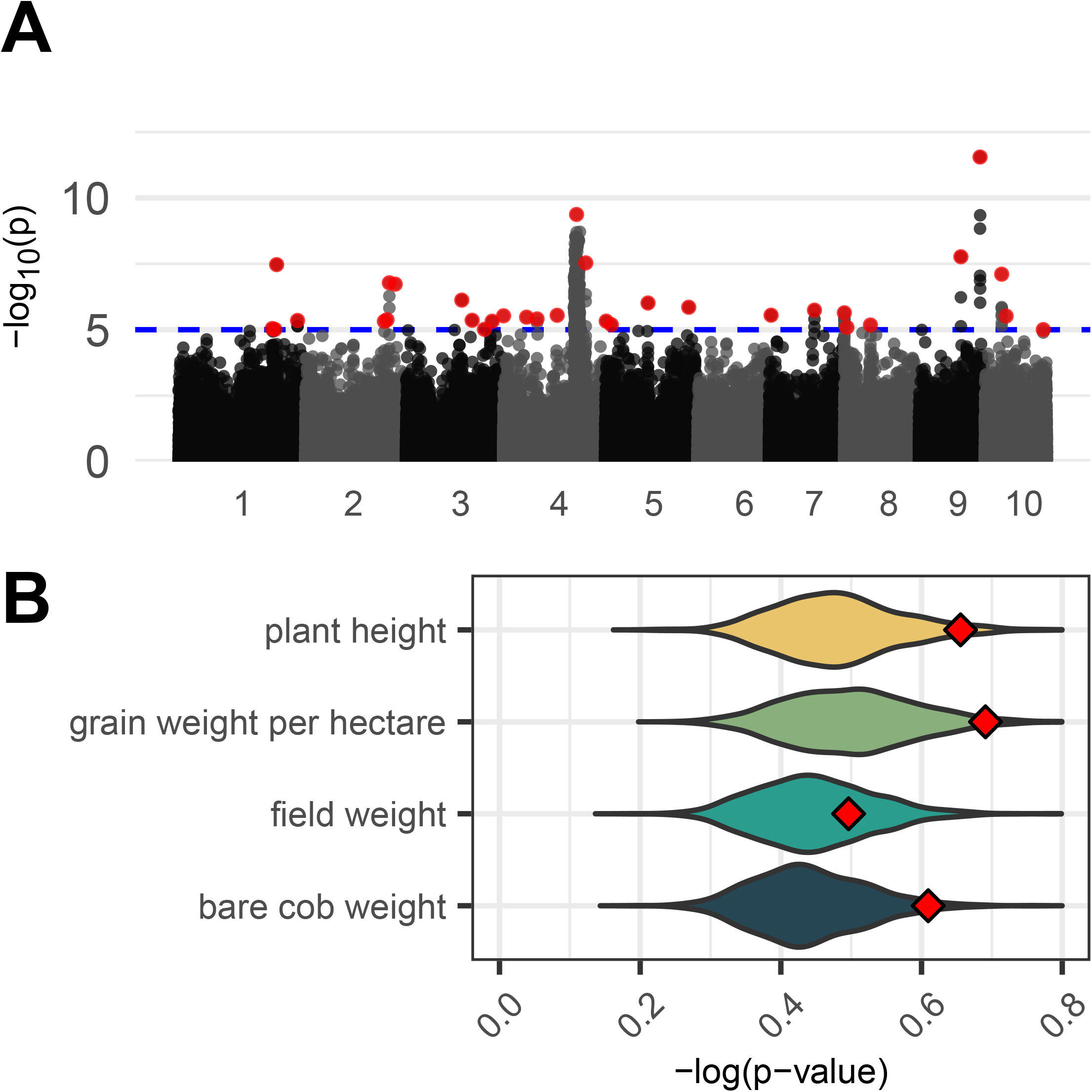
Multivariate envGWAS and phenotypic enrichment. **A)** Manhattan plot of multivariate envGWAS results testing each SNP for association against the set of seven climatic variables. Top SNPs representative of significantly associated loci in a nearby LD block are highlighted in red. Cutoff threshold of 10^*−*5^ represented by blue dashed line. **B)** Comparison of the average strength of phenotypic associations for the set of 32 top SNP hits selected by the envGWAS (red diamonds) against 1000 sets of 32 random SNPs of matching allele frequency (violin density plot). X-axis represents mean -log(p-value) of association to the given phenotypic trait for all SNPs in each set.

Although we find fewer loci than previous envGWAS of these data (Romero Navarro *et al*. 2017; Gates *et al*. 2019), our approach likely has a lower false positive rate (compare Supp. Figure S9 to Supp. Figure 14 in Romero Navarro *et al*. (2017)) due to our controlling for genetic relatedness among traditional varieties, and the way we accounted for multiple testing issues when running envGWAS on multiple correlated environmental variables. For example, we identify loci including *Inv4m* and a candidate SNP on chromosome 9 near an annotated heat-shock-related transcription factor *ZmHsftf9* (Zm00001d048041) (Gates *et al*. 2019). The *Arabidopsis* ortholog of this transcription factor (AtHSF1) is expressed under heat stress and is responsible for activation of heat-shock-protein-based thermotolerance (Schöffl *et al*. 1998; Lohmann *et al*. 2004), and the tomato homolog HSFA1 acts as a master regulator of thermotolerance (Mishra *et al*. 2002). In maize, ZmHsftf9 has been shown to have differential transcript splicing between high and low temperature regimes (Li *et al*. 2021) and is associated with yield differences under drought in African varieties (Gates *et al*. 2019).

### Are genotype-environment associations useful for breeding?

Given these phenotypic and genetic signals of local adaptation, we asked whether the loci that we discovered could help breeders select useful traditional varieties. As a first test of the value of envGWAS loci, we asked whether as a group these loci were significantly associated with phenotypic variation in our trials. Compared to random SNPs matched for minor allele frequency, envGWAS SNPs showed stronger associations to bare cob weight (*P* = 0.023), grain weight per hectare (*P* = 0.02), plant height (*P* = 0.037), and days to flowering (*P* = 0.02), but not for anthesis silking interval (*P* = 0.969) or field weight (*P* = 0.275) (Figure 4B).

To evaluate if these associated loci could explain phenotypic variation directly in genomic selection contexts, we estimated the percentage of variance in the yield trait BLUPs that could be predicted by these 32 loci (Figure 5). Since these loci may indirectly provide information on genome-wide population structure, we compared models with the top 32 envGWAS SNPs augmented by the first five principal components (PC) of the genomic relationship matrix (GRM) to a base model only using those PCs, as well as an alternative model including the PCs and random sets of SNPs matched by allele frequency with the envGWAS SNPs. The first two PCs were associated with elevation (Pearson’s correlation *r* = 0.83) and latitude of origin (*r* = *−*0.58) respectively (Supp. Table S6), but together explained less than 5% of the genotypic variance (Supp. Figure S1 and S2). We assessed predictive ability only within sets of testcross families sharing the same hybrid tester so that predictive abilities would not be inflated by differences in genetic values among testers. The model including just the first five principal components had the highest predictive ability across trials and tester for field weight (*r* = 0.305), followed by grain weight (*r* = 0.268), plant height (*r* = 0.207), and bare cob weight (*r* = 0.155) (Figure 5 and Supp. Figure S7). We found little evidence that including envGWAS SNPs in our models improved prediction accuracy for any of the yield traits beyond that captured by the genetic PCs (*P* = 0.999 for field weight, *P* = 1.00 for bare cob weight, *P* = 0.999 for grain weight, and *P* = 0.997 for plant height), and little evidence that they improved prediction over randomly selected SNPs (*P* = 0.999 for field weight, *P* = 0.999 for bare cob weight, *P* = 0.9963 for grain weight, and *P* = 1.00 for plant height).

**Figure 5.**
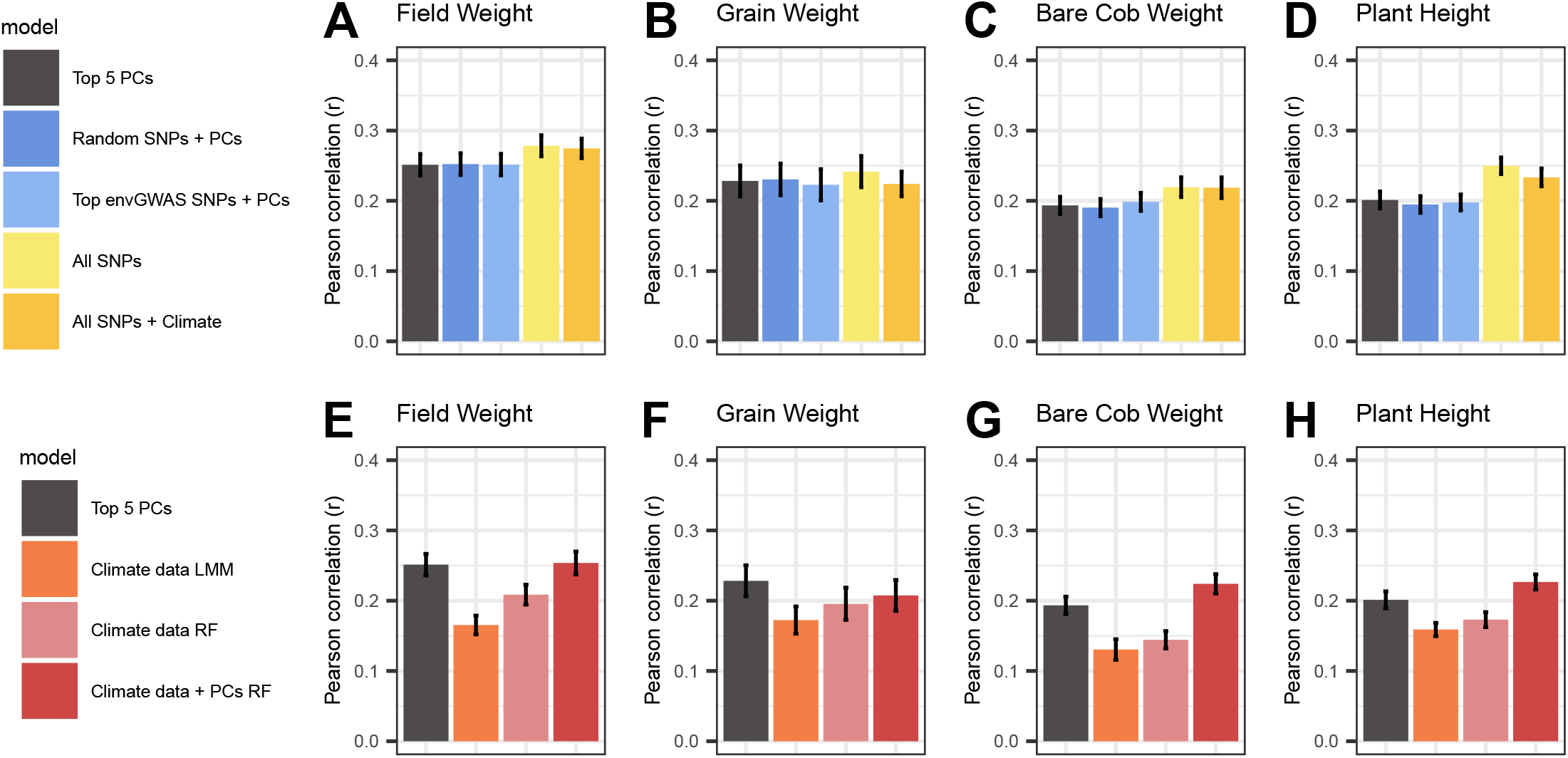
Genomic and environmental prediction models. Comparison of prediction models predicting field weight, grain weight, bare cob weight, and plant height. Y-axis represents predictive ability (Pearson’s *r*) between a traditional variety’s predicted value and actual BLUP value. Bar plots represent mean predictive ability across all tester families, and averaged across all trials, colored by selected genomic models **(A-D)** and environmental models **(B-H)**.

We compared these models to two alternative strategies for predicting the value of traditional variety alleles for increasing yield. We fit genomic prediction models using all 300K GBS SNPs as well as environment-of-origin prediction models using the data associated with each traditional variety (similar to FIGS models). The first model evaluated the total value of all (common) alleles from the GBS data irrespective of their environmental distribution. The second model evaluates the gain in predictive ability provided by environmental data beyond that already captured by genetic information.

The genomic prediction model using all SNPs explained significantly more phenotypic variation in field weight than the PC-only (*P* = 0.045) and envGWAS SNPs + PC models (*P* = 0.013). We also found the all-SNPs model to have higher predictive ability for bare cob weight (*P* = 0.001 and *P* = 0.001 relative to PC-only and envGWAS SNPs + PC, respectively) and plant height (*P* < 0.001 and *P* < 0.001, respectively), but the improvement was not significant for corrected grain weight (*P* = 0.228 against the PC-only model and *P* = 0.098 against the envGWAS SNPs model) In contrast, environment-of-origin prediction models that included PC features did not significantly increase predictive ability over the PC-only model (*P* = 0.999 for field weight, *P* = 0.997 for bare cob weight, *P* = 1.000 for grain weight, *P* = 0.998 for plant height), and the environment-of-origin prediction models that included all SNPs did not significantly increase predictive ability over the genomic prediction model (*P* = 0.999 for field weight, *P* = 0.997 for bare cob weight, *P* = 1.000 for grain weight, *P* = 0.998 for plant height).

These results consistently show that environment-of-origin data provides little additional benefit for predicting variation in yield traits among traditional varieties beyond that captured by GBS markers. However, environment-of-origin data is still more accessible than GBS data for many germplasm collections. To test whether environment-of-origin are useful in this dataset when GBS data are not available, we assessed the accuracy of predictive models using only environment-of-origin data in linear models and random forests (Figure 5B and Supp. Figure S8). Predictive abilities of these environment-of-origin only models were significantly greater than zero on average (*P* < 0.001 for all traits), but generally had low accuracy, regardless of the methodology chosen.

## Discussion

Today’s plant breeding programs need to develop crop varieties that will be resilient to the effects of climate change and provide high yields in future climates. We hypothesized that existing germplasm collections of traditional maize varieties, collected across a wide range of climate gradients, carry useful diversity for adapting existing varieties to new environments. Using a large panel of geo-referenced collections with high-density genotyping, we evaluated whether environmental data from these collections could be used to prioritize traditional varieties for breeding. Despite the fact that these accessions exhibit both genomic and phenotypic evidence of local adaptation, we found that incorporating environmental data from their collection locations provided little additional information directly in genomic breeding.

On the one hand, our results are encouraging in that we find strong evidence of local adaptation. From our field trial, we show that environments-of-origin of traditional varieties are correlated to yield traits (Figure 2) and thus should be informative when considering agronomic performance across diverse environments. We find this despite a likely attenuated local adaptation signal in our study design, as traditional varieties were crossed mostly to hybrid testers adapted to similar elevations, and were mostly grown at these elevations. This was done to more closely reflect the breeding process at those elevations, as maize has been shown to exhibit asymmetrical local adaptation, where individuals from highland environments have lower fitness in lower altitudes whereas lowland-origin individuals perform equally well across altitudinal clines (Samayoa L. *et al*. 2018; Mercer *et al*. 2008). Likewise, even after accounting for spatial autocorrelation and population structure, our GPoE analyses find meaningful correlations between predicted and observed environments, pointing to a genotypic signal of local adaptation. Consistent with previous analyses of these data (Gates *et al*. 2019), we identify a number of convincing outlier loci, including *Inv4m*, a large inversion introgressed from *Zea mays spp. mexicana* (Pyhäjärvi *et al*. 2013; Hufford *et al*. 2013) that appears to impact flowering time (Romero Navarro *et al*. 2017), and the heat shock protein *hsftf9* that shows differential splicing under different heat regimes (Li *et al*. 2021). Combined, the envGWAS-discovered loci are enriched for associations with phenotypes of agronomic importance (Figure 4) indicating that envGWAS successfully identifies loci of functional relevance.

On the other hand, while we can identify environment-associated alleles from envGWAS that are enriched for phenotypic relevance, our results suggest that the specific loci that we discovered by envGWAS do not directly improve predictions of phenotypic values of traditional maize testcrosses in modern agronomic contexts and environmental characterization of traditional maize origins appears only weakly predictive of genetic values for breeding.

### The potential and limitations of envGWAS

envGWAS is an appealing technique for discovering candidate loci driving local adaptation to environment gradients because it can be applied to large collections of traditional germplasm without the need for expensive field trials. However, like any GWAS method, envGWAS cannot be comprehensive – it will likely miss many of the most valuable alleles (false negatives) and can also be subject to high false positive rates.

There are several possible reasons why envGWAS may miss important loci relevant for breeding. First, GWAS prioritizes variants based on the amount of variance in the trait of interest, while breeders are most interested in alleles with large effect sizes. The amount of variance explained by a locus is proportional to its allele frequency. Because large effect alleles will be kept rare under purifying or stabilizing selection (Pritchard and Cox 2002), envGWAS may miss many loci of large effect.

Second, because LD decays quickly in traditional maize varieties (Chia *et al*. 2012), the ability of GWAS approaches to identify causal genes or QTL from sparse sequencing like GBS is impacted (Tiffin and Ross-Ibarra 2014). The *≈* 350, 000 GBS markers in our dataset are nonrandom and constitute a small fraction of the common variants in maize (Andorf *et al*. 2024). Higher-density genotyping of this population would likely help discover more causal loci. Finally, even if the loci driving local adaptation in these diverse panels had sufficiently large effects, sufficiently high allele frequency, and were in LD with GBS markers such as to be detectable by envGWAS, adaptation may be controlled by latent environmental variables that we did not measure or are otherwise unknown. We included seven environmental variables describing variation in climate relevant to plant growth and development in specific growing seasons. However, there are many other environmental and management variables that affect plant performance that we did not include, such as soil variables, planting densities, and biotic pressures (Costa-Neto *et al*. 2021). Alleles that increase performance along these gradients may have contributed to local adaptation or be useful in certain agronomic contexts but would have been missed by our analysis.

At the same time, our envGWAS may also be sensitive to mechanisms that create false-positive associations that do not control yield in modern agronomic contexts. While false positives may be common due to poorly calibrated statistical tests that run envGWAS independently for each variable, we use MegaLMM and JointGWAS to de-correlate genotype-environment and environment-environment relationships, resulting in well-calibrated statistical tests which should reduce our false positive rate.

Nonetheless, loci that controlled yield in traditional agronomic contexts, but do not control yield in our target environments could show up as false positives because they are not useful for future improvements in yield. We might find such loci if they controlled responses to environmental variables that were correlated with any of our seven focal environmental variables across traditional farms. For instance, since biotic stresses such as disease are correlated with rainfall or humidity (Cairns *et al*. 2012), loci underlying disease resistance may be discovered as having an association with precipitation level, despite the causal relationship being disease pressure on the locus. However, since disease pressure was limited by pesticide applications in many of our trials, such loci may not have impacted the yields that we observe.

Finally, population structure and spatial autocorrelation of environmental variables can generate non-adaptive associations between neutral alleles and environmental variables Lasky *et al*. (2023); Blanquart *et al*. (2012);Günther and Coop(2013). Population structure exists within maize traditional varieties for many reasons including dispersal patterns shaped by geographic distance, human migration, farmers’ preference for quality traits, seed sharing via trade, and introgression of wild relative alleles in limited geographic regions (Kennett *et al*. 2022; Louette and Smale 2000; Bellon *et al*. 2003; van Heerwaarden *et al*. 2011). If populations also occupy different environmental zones, those divergent alleles will also associate with environmental gradients.

We attempted to reduce the number of such false positive discoveries using an estimated genomic relationship matrix as a covariate in our envGWAS, which greatly reduced the number of significant discoveries relative to earlier analyses of this dataset (Romero Navarro *et al*. 2017; Gates *et al*. 2019). However, this correction will also restrict the discovery of any alleles that drive population differentiation, including true positive adaptation to those environmental gradients. To mitigate this effect, we used a leave-one-chromosome-out (LOCO) strategy in calculating the GRM, which strikes a balance between sensitivity (reducing false negatives) and specificity (reducing false positives) due to population structure (Yang *et al*. 2014). Future efforts that model genetic background and migration by accounting for spatial effects (Lasky *et al*. 2024) may improve the true discovery rate of envGWAS.353

### Evaluation of alternative strategies for discovering useful traditional varieties

Genome scans such as envGWAS are only one plausible way to identify useful genetic diversity in germplasm collections. Rather than using georeferencing data and genomic data to identify specific candidate loci, the same data can be used to identify varieties that likely perform better in distinct environments through genomic prediction and incorporation of environmental predictors. In this study, we compared the effectiveness of using environment-of-origin data to the value of envGWAS-discovered alleles in phenotypic studies.

We found that environment-of-origin data of traditional varieties modeled alone explained a small but significant percentage of the variation in yield (Figure 5). This finding corresponds with our GPoE analyses showing strong genetic differentiation of varieties across environmental gradients. This is also externally consistent with extensive previous evidence that maize traditional varieties are locally adapted (Janzen *et al*. 2022), as well as previous analysis of a subset of Mexican accessions in this dataset by McLaughlin *et al*. (2024) showing that random forest models utilizing only environment-of-origin data were capable of predicting some root anatomy traits. If genotyping every individual in a diversity panel is prohibitively expensive or population structure is unknown, environment-of-origin data of collections may thus enable selection of a starting pre-breeding population.

However, our study shows that environmental data explained negligible additional variation in field trials beyond what was predictable using genomic data. Population structure, isolation-by-distance, and spatial autocorrelation in environment can generate correlations between environmental values and performance that would be equally captured by genomic data (Bahn *et al*. 2008; Blanquart *et al*. 2012). Evidence of this is seen in our GPoE results, which show that the association between genotype and environment is diminished once accounting for spatial proximity, a proxy for population structure.

In contrast to predictions derived from environmental data or environmentally-associated SNPs, models using whole-genome GBS markers had reasonably high predictive ability for traditional variety crosses. This suggests that given existing field trial data in relevant environments, yield predictions from genome-wide genotyping data likely capture more dimensions of environmental adaptation and selection than would be captured from environmental variables selected *a priori*. However, our results suggest much of the value of genome-wide genotyping may simply be the quantification of population structure. The GBLUP prediction model we used gains accuracy from both historical LD with causal markers and from linkage due to population structure (Habier *et al*. 2007,2013). We find that the first 5 PCs of the genotyping data provide predictive ability similar to the full kinship matrix, suggesting that much of the predictive ability of GBLUP comes from population structure. While this has been previously shown to be true in established breeding populations of hybrid temperate maize (Windhausen *et al*. 2012), we show that this occurs in diverse traditional variety collections as well. Learning from which populations a traditional variety is collected from may provide information to help breeders prioritize which germplasm to introduce in their programs. However, this is a relatively coarse type of information dependent on the population of interest, and may not help breeders leverage the full value of a large germplasm collection.

### Utility of environmental data for pre-breeding programs

Unlike Lasky *et al*. (2015), who found that incorporating the top 100-250 representative envGWAS SNPs outperforms kinship-only GBLUP models in panicle weight prediction, our work did not find a significant change in predicting yield component traits using a limited number of envGWAS SNPs in a GBLUP model. Our results were in concordance with a previous analysis of this maize dataset for flowering time by Romero Navarro *et al*. (2017), who found similar predictive ability for flowering time using genome-wide SNPs and a large set of envGWAS markers. In addition, our results were similar to those of Kehel *et al*. (2020) who showed that models including passport environmental data of traditional varieties do not provide a significantly higher predictive ability than models with only genomic data, even with higher SNP density.

Despite not being useful for genetic value predictions in our collections, our results don’t preclude envGWAS-identified loci being used in other ways in germplasm collection characterization and prebreeding. For example, individual envGWAS may still identify functionally relevant alleles that could be incorporated into breeding via marker-assisted selection or genome editing (e.g. Barnes *et al*. 2022). Additionally, envGWAS loci could be used in breeding by first selecting high-yielding traditional varieties in target environments via genomic prediction, and then ranking the selected varieties based on the enrichment of novel envGWAS loci not already present in elite hybrids. Subsequent F2 mapping populations could then be used to evaluate adaptive envGWAS loci in Origin-Specific Genomic Selection (OSGS), partitioning performance based on parental origin (Yang *et al*. 2020). Finally, environmental data may continue to be useful for predicting gene-environment interactions where differences in environmental conditions between trial locations can be leveraged to identify gene-by-environment effects and select varieties with favorable responses to specific environmental stresses (Lasky *et al*. 2015; Costa-Neto *et al*. 2021).

## Materials and Methods

### Samples and Genotyping

The Seeds of Discovery project (SeeD) has characterized the CIMMYT International Germplasm Bank maize collection containing over 24,000 accessions of traditional varieties primarily from North, Central, and South America. Here, we used data generated across a core collection of *≈* 4, 000 accessions from this project representing the breadth of environmental variation across the Americas as described in (Romero Navarro *et al*. 2017). Accessions were narrowed down to 3511 genotypes representing 2895 accessions based on availability of accurate coordinates for each accession’s collection location, where 6-month growing season environment data was complete for all variables of interest (see details in **Environmental data** below).

Briefly, each individual was sequenced using genotyping-by-sequencing (Elshire *et al*. 2011) to a median 2X coverage. Genotypes were called using TASSEL (Glaubitz *et al*. 2014), and imputed using BEAGLE 4 (Browning and Browning 2013) resulting in an initial set of 955,120 imputed SNPs reported in (Romero Navarro *et al*. 2017) using the B73 v4 reference genome (Jiao *et al*. 2017). Genotypes with more than 25% missing data and biallelic SNPs with minor allele frequency > 0.01 were kept after filtering, resulting in a final filtered set of 345,270 SNPs. To estimate kinship, a genomic relationship matrix 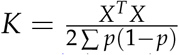 was estimated using SNPs above MAF > 0.05 for the genotype matrix *X* as per Goudet *et al*. (2018).

PCA analysis was performed using the R package *prcomp* on a subset of 5000 randomly sampled SNPs taken genome-wide from the filtered and imputed GBS data.

### Environmental data

For each accession in our dataset, environment data was obtained using the geographic coordinates attached to collection metadata. Monthly estimates of minimum and maximum temperature and total precipitation were obtained from WorldClim (Fick and Hijmans 2017), aridity measurements were obtained from the Global Aridity Index and Potential Evapotranspiration Database (Zomer *et al*. 2022), relative humidity measurements were obtained from ERA5 (Bell *et al*. 2021), and elevation for each accession was taken from the SRTM 90m Digital Elevation Database v4.1 (Jarvis *et al*. 2008). Environmental variables were calculated across 6-month growing seasons for each accession as defined by the growing agroecological zone from the FAO (Fischer *et al*. 2021).

Variables calculated included: minimum temperature (tmin) over the growing season; maximum temperature (tmax) over the growing season; range of temperature (trange) as defined by tmax *−* tmin; total precipitation (precipTot) accumulated over the growing season; mean aridity (aridity Mean); mean relative humidity (rhMean), and elevation at point of collection for each accession. We include max temperature and minimum temperature as environmental variables as it is known that heat stress affects developmental and physiological traits such as photosynthesis and shortened life cycle (Muchow *et al*. 1990; Crafts-Brandner and Salvucci 2002). Likewise, as precipitation levels such as high rainfall and drought affect maize yields (Li *et al*. 2019), we include total precipitation for our model. As vapor pressure deficit is a major crop model covariate driving yield (Seetharam *et al*. 2021), we include relative humidity and mean aridity measures as an indirect proxy measure. Supp. Figure S3 illustrates the distributions and correlations between environmental variables covered by this population. To model non-Gaussian distributions of environmental variables such as precipitation in a non-parametric manner, we calculated the ranked inverse normal transformation (INT) for the whole population using the formula

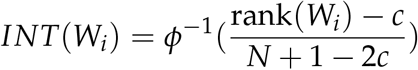

where we transform each environment value *W*_*i*_ using the conventional Blom’s offset *c* = 3/8 (McCaw *et al*. 2020; Tang and Lin 2015).460

### Trials

As part of the SeeD project evaluation of the traditional variety core collection, offspring of traditional variety collections and groups were planted in multiple environments under a replicated F1 crossing design known as F-One Associated Mapping (FOAM) (see Romero Navarro *et al*. (2017) for design details). Here, we analyze data on plant height (PH), the total weight of ears (kernels and cob) measured in the field (field weight (FW)), bare cob weight (BCW), and grain weight per hectare adjusted at 12.5% of humidity (GWPH) from (Gates *et al*. 2019), as well as flowering traits (days to female flowering (DtF), and anthesis-silking interval (ASI)) from (Romero Navarro *et al*. 2017).

Two important features of the crossing experiment were included to ensure that phenotype data was not overly biased by elevational adaptation. First, plants were preferentially grown in locations that were of similar adaptation (highland tropical, sub-tropical or lowland tropical) to their home environment. While F1s with a highland accession parent were grown in low elevation and vice versa, on average, more plants from highland accessions were grown in high elevation than low elevation trials. Second, each plant was crossed to a tester that was adapted to the environment that the F1 seeds were grown in, for a total of *≈* 4700 testcross genotypes (Supp. Table S4 shows the number of testers used in each trial). Both of these design features facilitate comparison of a larger sample of accessions, but also lead to an unbalanced experimental design and reduce apparent adaptive differences among traditional varieties, making our estimates of adaptation more conservative.

Briefly, the F1 offspring of each accession were grown in ambient field conditions across a total of 23 trials in 13 locations in 2 years. Each testcross family was grown in a single plot with a range of 9-26 (average of 16) offspring per accession, with an average of 850 accessions (collections and groups from the seed bank) grown in each trial in an augmented row-column design. As we were only interested in accessions with available geographic coordinate data, we considered only accessions that were labeled as collections: Supp. Table S3 shows the number of collection accessions with environmental data measured for each phenotypic trait in each trial, averaging 586 per trial. While this range of field environments (Supp. Table S1) is exceptional with respect to the standard for field based studies attempting to identify the genetic basis of environmental adaptation, we do note that there are a number of maize environments (e.g. extreme high elevation above 2500m, tropical with invariant daylength, and wet, low elevation) that may not be well represented here but should be of interest to future studies of this nature. Across the 23 trials, plants were phenotyped for a variety of agronomically relevant traits (Supp. Figure S2). In summary, our field trials included 17,902 plots spanning trial locations, testcrosses, and accessions, with 76,042 phenotypic observations.

### Phenotypic data

Breeding values for each traditional variety were estimated as described in Navarro *et. al*.(Romero Navarro *et al*. 2017), controlling for design variables in a complete nested model. Briefly, raw phenotype values were converted into best linear unbiased predictions (BLUPs) that controlled for trial, checks, field position, plot, and tester effects.

To ensure that effect sizes of genetic values have appropriate scale across trials and to correct for shrinkage, BLUPs were deregressed (Garrick *et al*. 2009). Trait heritability scores for each trial were calculated as 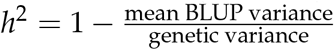. For each trait analyzed, we kept only trials with heritability scores greater than 0.1 for that trait, and removed trial:trait:tester combinations with less than ten unique values to filter out populations with low sample sizes. In total, 2689 genotypes representing 2518 unique collection accessions with sufficient sample size across trials and had environmental data connected to collection coordinates were used for phenotypic analyses.

### Local adaptation

Similarly to Gates *et al*. (2019), we tested for evidence of local adaptation by estimating the relationship between the environment-of-origin of each traditional variety and the yield trait values of the testcross families in the field trials. Local adaptation is indicated if traditional varieties collected from environments more different from the environment of each trial have lower values of the yield traits (Wadgymar *et al*. 2022).

To test for this, we fit a statistical model relating the environment-of-origin and the environment of each trial to the variation in yield trait values in each trial. Here, we used elevation, mean temperature, and annual precipitation measurements from WorldClim 2 (Fick and Hijmans 2017) so that time was consistent between maize collection and trial location coordinates. We fit a quadratic function to trait values within each trial as a function of the environment-of-origin value of each traditional variety, and modeled the environmental value that maximized this function as a function of the environment of the trial. Specifically, we fit:

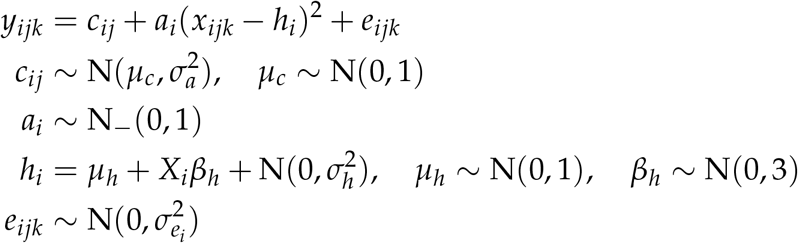

We fit this model separately for each trait and each candidate environmental variable, where *y*_*ijk*_ is a z-scaled de-regressed BLUP of F1 testcross family *k* crossed to tester *j* in trial *i* with climate value of the accession *x*_*ijk*_ and climate value of the trial *X*_*i*_. *c*_*ij*_ represents the maximum value of each response function, unique for each tester for each trial, *a*_*i*_ is the quadratic parameter specifying the curvature of the response function and constrained to be non-positive through its prior, and *h*_*i*_ is the coordinate of the maximum of the response function in each trial. We modeled *h*_*i*_ as unique for each trial, but with a possible relationship to the climate value of the trial *X*_*i*_, with regression coefficient *β*_*h*_. If *β*_*h*_ is positive, it implies that trials with higher values on a climate axis select for accessions originating from high climate values. If *β*_*h*_ = 1 and *µ*_*h*_ = 0, accessions from similar climates to the trial had highest fitness values. Therefore, positive values of *β*_*h*_ indicate local adaptation, while larger (negative) values of *a*_*i*_ indicate stronger decreases of fitness as accessions’ environment-of-origin deviates further from the optimum *β*_*h*_. We implemented this model using the R package *brms* to obtain posterior values for these parameters (Bürkner 2017).

### Genomic Prediction of Environment (GPoE)

GPoE analyses to predict environmental variables from genotypic data (*E ∼ G*) was performed with MegaLMM (Runcie *et al*. 2021). As no phenotypic data was necessary for this analysis, all 3511 genotypes that had both available genomic data and associated environmental data were used. Briefly, MegaLMM decomposes the covariance of genetic random effects and environmental variables simultaneously to de-correlate environmental relationships and genetic relationships, and uses those covariance matrices to estimate withheld testing data. The kinship matrix *K* was calculated as the genomic relationship matrix above and served as the input for genotype data. The model for prediction was **Y** = **FΛ** + **U**_*R*_ + **E**_*R*_ where **Y** represents a matrix of environmental values for 3511 genotypes and seven INT-transformed environmental variables, **F** and **Λ** are factor scores and loadings for *k* = 7 latent factors, and **U**_*R*_ and **E**_*R*_ are matrices of residual genetic and non-genetic values not explained by the factors. **F** was also modeled as a function of genetic and non-genetic effects: **F** = **U**_*F*_ + **E**_*F*_. Matrices **U**_*R*_ and **U**_*F*_ were matrix-normal random variables with diagonal column covariances and row covariances proportional to **K**. Matrices **E**_*R*_ and **E**_*F*_ were matrix-normal random variables with diagonal row and column covariances. MegaLMM was run with default parameters (Runcie *et al*. 2021) with a burn-in of 2000 iterations, and convergence reached over another 2000 iterations.

K-fold cross-validation was performed for *k* = 10 folds, with a testing set of approximately 350 accessions having environmental data withheld before training. Training was done on non-withheld data through MCMC sampling via MegaLMM: 200 samples were collected each iteration. After 20 iterations of sampling, predicted **Û** values for each environmental variable was calculated as the posterior mean for both testing and training set accessions: **Û** = **U**_**R**_ + **U**_**F**_**Λ** To evaluate predictive ability, Pearson’s *r*^2^ correlation for each environmental variable was calculated as **cor**(**Û**, **Y**)

To test the effect of geographic distance on genotype-environment correlations, we used the R package *spatial sample* (Mahoney *et al*. 2023) to split our study population into ten geographically separated folds, clustered non-deterministically based on distance. Cross-validation was then performed as described above for our *k* = 10 spatial folds, withholding accessions for a given fold with a buffer distance of 400 km to prevent accessions directly bordering the fold to inform prediction for the accessions within the fold, thereby enforcing spatial independence. A pairwise t-test was performed to test whether a significant difference between random and spatial k-fold cross-validation existed for each environmental variable and trial combination.

### Environmental GWAS (envGWAS)

Environmental genome-wide associations (envGWAS) were run for each environmental variable (elevation, tmax, tmin, trange, rhMean, precipTot, aridityMean). The same 3511 genotypes that had both genomic data and associated environmental data as in the GPoE analysis were used for envGWAS. Associations were made between the filtered SNP set and INT-transformed growing season data described above.

envGWAS was performed using the R package *JointGWAS* (Runcie 2022). JointGWAS tests each SNP for association with variation in *any* environmental variable using an F-test. Residual correlations among environmental variables and correlated genetic background effects across genotypes were accounted for using genetic and residual covariance matrices estimated using MegaLMM (Runcie *et al*. 2021). Markov-Chain Monte Carlo sampling to obtain variance estimates for each variable was performed with a burn-in of 2000 iterations, and convergence reached over another 2000 iterations.

The association model was run in JointGWAS for each SNP as follows:

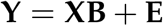

where ***Y*** was the ***n*** × ***t*** matrix of environmental values, ***X*** was the ***n***× ***b*** genotype design matrix, and ***B*** was the ***b*** × ***t*** matrix of estimated genotype effects.

**E** was modeled as a random variable with multivariate normal distribution with mean zero and covariance **Z**(**G** ⊗ **K** + **R** ⊗ **I**)**Z**^*T*^, where **G** and **R** are background genetic and residual covariance matrices among environmental variables estimated as the posterior means of these matrices from Mega LMM run with default parameters, **K** is the genomic relationship matrix computed above, **I** an identity matrix, and **Z** represented the design matrix relating all accession : environmental value combinations to the observed environmental values. For computational speed, Joint GWAS pre-multiplies both sides of the association model by the inverse of the Cholesky combination of the residual covariance matrix, turning the model into a standard linear model, after which a normal F-test was applied to the coefficient of the **X** term.

To account for population structure and inter-chromosomal LD, the genomic relationship matrix **K** was separately calculated for each chromosome and the association test run separately per chromosome as well. A p-value significance threshold of 10^*−*5^ was used to identify SNPs that were considered significant to any environmental variable, set at a slightly lower threshold than the Bonferroni correction to evaluate more loci for phenotypic enrichment and prediction i.e. for pre-breeding introgression.

### Phenotypic association

In addition, phenotypic genome-wide associations were run for each trait (PH, FW, BCW, GWPH, DtF, ASI) across all trials using JointGWAS. Here, we test whether each SNP has association with phenotypic variation in *any* trial with an F-test.

As in the envGWAS, genetic and residual covariance matrices were estimated using MegaLMM, but accounting for residual correlations among trials and genetic background. Since we measured the phenotypes between F1 offspring of our accessions, we divide our genomic relationship matrix *K* by a factor of 4 to account for half-sib kinship. The same default parameters for MegaLMM were used as in the envGWAS.

The association model for each SNP was:

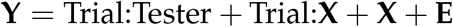

where **Y** was the matrix of BLUP values over all trials, Trial:Tester were fixed effects, **X** was the main effect of a SNP, Trial:**X** was the interaction between each SNP and each trial. **E** was modeled the same as in the envGWAS above, but where **G** and **R** are background genetic and residual covariance matrices among trials from MegaLMM, and **Z** a design matrix relating all possible accession:trial combinations (columns) to the observed combinations (rows). To increase GWAS power, when testing each SNP we excluded all SNPs on the same chromosome when calculating **K** for the covariance computation. The genome-wide **K** was used with MegaLMM when estimating **G** and **R**.

Finally, normal F-tests were applied to the coefficients of both the terms **X** and Trial:**X**. The p-values from these two F tests were combined using Fisher’s method into a single test of any association of the marker in any of the trials.609

### Enrichment of phenotypic consequence for envGWAS SNPs

From our envGWAS, we take the top SNPs most significant for environment and observe if those SNPs similarly had significantly higher effect on yield or developmental traits from our phenotypic trials. To prevent over-representation from envGWAS peaks with large genomic regions encompassing many adaptive loci such as *Inv4m*, we opted to measure the effect of a given representative SNP obtained through LD clumping. Clumping was performed genome-wide to capture trans-LD across chromosomes as well as to identify mis-mapping in the assembly. LD clumping was performed by calculating the pairwise *r*^2^ of all SNPs above the 10^*−*5^ p-value significance threshold. We wrote a custom Python script to perform clumping using a greedy algorithm, prioritizing the most significant SNPs as lead SNPs and assigning SNPs to a lead SNP if they had correlation *r*^2^ > 0.3 to a lead SNP. Under these parameters, we obtained 32 lead SNPs representing top peaks in our envGWAS results.

Enrichment analysis of our envGWAS SNPs in our phenotypic GWAS was performed by taking our top 32 SNPs and obtaining 1000 samples of 32 random SNPs across the genome with minor allele frequency (MAF) matching the distribution of that of our envGWAS SNPs. For each phenotypic trait, we pulled the *−log*(p-value) from the phenotypic GWAS for both our lead envGWAS SNPs and each set of matching MAF SNPs. We then compared whether the average *−log*(p-value) for the envGWAS SNPs was an outlier compared to the null distribution of randomly sampled SNPs. P-values were calculated by finding the fraction of matching SNP sets with a larger average *−log*(p-value) compared to the envGWAS SNP set.

### Candidate Genes

To identify candidate genes from our envGWAS, we use the 32 lead SNPs obtained above from LD clumping, such that SNPs significant for environment are grouped together with SNPs within LD. Candidate genes were then chosen based on nearest genomic distance to lead clumped SNPs using BEDtools (Quinlan and Hall 2010) against the gene model annotation of the V4 assembly of the maize genome (Jiao *et al*. 2017). We then filtered down genes to only that had curated annotations and known models in the V4 assembly of the B73 reference genome. LD analysis was performed by comparing *r*^2^ of all SNPs in the GBS sequencing within a 300kbp upstream and downstream range of a given lead SNP, with the exception of S4_177835031, in which we used the 600kbp range.

### Genomic prediction (GP)

We performed genomic prediction on the deregressed phenotypic BLUP data, using the accessions with available climate data. We predicted the genomic BLUP (gBLUP) value for each accession in our panel and compared it to the observed BLUP value from field trials for each phenotypic trait.

As each testcross’ performance is partially controlled by the tester’s genotype, which was non-randomly assigned with respect to both traditional variety genotype and environment-of-origin, we estimated the predictive ability of each model by calculating the Pearson’s correlation between predicted and observed values within sets of testcrosses made with the same tester. We did this using cross-validation by assigning all testcrosses made with the same tester to the same fold, and trained models on yield BLUP residuals (corrected for tester ID) from all other testcrosses. Models were fit and evaluated separately for each trial

Linear prediction models were constructed in rrBLUP (Endelman 2011) with the general model *Y* = *Xβ* + *Zu* + *ϵ* where *Y* represented the BLUP residuals of each sampled individual, correcting for tester ID. For all models, a design matrix *X* of tester genotypes remaining in the training set was included as a fixed effect. Principal components were calculated using the singular value decomposition of the kinship matrix for the accessions present in each trial. The first five eigenvectors were included as features in each model.

We constructed the following models:

1. A baseline PCs-only model where *X* included tester and the first five principal components.
2. A PCs + envGWAS SNPs model where we added a 3511 × 32 random effect matrix *Z* of the lead SNPs representing the 32 loci significantly associated with environment from our envGWAS analysis.
3. A PCs + random SNPs model where we additionally created an equivalent 3511 × 32 random effect matrix *Z* matrix of randomly sampled SNPs from the genome with matching allele frequency to the envGWAS SNPs.
4. An all-SNPs model where the random effect *u* had covariance *K*, the kinship matrix calculated from all SNP markers in the GBS data, in addition to the *X* fixed effects matrix of tester and first five PCs.
5. An all-SNPs + environment model that was identical to the previous, but also included the INT-transformed environment-of-origin data for each accession as fixed effects in the *X* design matrix. Further, to assess how how well phenotypic variation could be predicted by environmental data, we additionally tested models given only aggregate environment data, and with models employing non-parametric random forest models using the R package Random Forest (Liaw and Wiener 2002) with ntree = 1000. Random forests were run with 1000 decision trees. These included:
6. A linear model in rrBLUP with a *X* matrix with only tester + climatic data;
7. A random forest with tester + climatic data as predictor variables;
8. A random forest with tester + climatic data + first five PCs as predictor variables.

To measure differences in predictive ability between the first five models, we fit the following linear mixed model to the correlation measures of each model in each fold using the *R* package *lme4* (Bates *et al*. 2015):

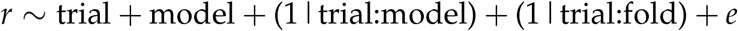

where the formalism (1|X) represents a random effect with grouping level specified by the factor X. Since the sampling variance of Pearson’s r is higher for smaller sample sizes, residuals *e* were weighted by the inverse of the sample size. Comparisons among the main effects of the models were evaluated using Tukey’s method implemented in the *emmeans* and *lmerTest* packages (Searle *et al*. 1980; Kuznetsova *et al*. 2017).

## Data Availability and Benefit-sharing

The Github repo with analysis scripts is available at github.com/liforrest/SeeD. Imputed GBS SNP data used for genetic analysis is available at https://data.cimmyt.org/dataset.xhtml?persistentId=hdl:11529/8702394. Curated information of field trial metadata and phenotypic BLUP data is available at https://data.cimmyt.org/dataset.xhtml?persistentId=hdl:11529/10548233 This research collaboration was made possible through the CIMMYT Seeds of Discovery project. All collaborators are included as co-authors, and benefits from this research are made available to shareholders and the public community as per the CIMMYT Standard Material Transfer Agreement (https://www.cimmyt.org/content/uploads/2019/04/SMTAFAQCIMMYTSept09.pdf) Our research is committed to the equitable sharing of knowledge and benefits with institutions and peoples who have contributed to this work.

## Acknowledgments

FL would like to acknowledge funding from the Gates Foundation and the Jastro-Shields research award. Additionally, the authors would like to acknowledge funding from the NSF: award number 1546719 to JRI and award number 1238014 to ESB. ESB also acknowledges support of the USDA-ARS, JRI acknowledges support from USDA Hatch project CA-D-PLS-2066-H, and RRA acknowledges support from USDA Hatch Project 02776. CIMMYT would like to acknowledge the Sustainable Modernization of Traditional Agriculture (MasAgro) project supported by the Ministry of Agriculture and Rural Development (SADER) of the Government of Mexico for funding the Seeds of Discovery maize traditional variety characterization. Finally, we acknowledge the smallholder farmers and indigenous people whose work and love for their traditions and identity keep maize diversity alive.

## Notes

### Competing Interest Statement

The authors have declared no competing interest.

