## Supplemental Figures and Tables for "The utility of environmental data from traditional varieties for climate-adaptive maize breeding"

### Supplementary Tables and Figures

| Trial ID | Year | Location | Elevation (m) | Trial latitude | Trial longitude |
|---|---|---|---|---|---|
| m2011BAFNN | 2011 | Agua Fria | 430 | 20.46 | -97.46 |
| m2011BCH | 2011 | G Victoria | 800 | 16.44 | -93.13 |
| m2011BCL | 2011 | Celaya | 1894 | 20.58 | -100.82 |
| m2011BMO | 2011 | Tarimbaro | 2073 | 19.78 | -101.18 |
| m2011BNY | 2011 | San Pedro | 1424 | 21.22 | -104.73 |
| m2011BTLNN | 2011 | Tlaltizapan | 905 | 18.69 | -99.13 |
| m2011BTORN | 2011 | Torreon | 1137 | 25.56 | -103.37 |
| m2012AAFCA | 2012 | Agua Fria | 430 | 20.46 | -97.46 |
| m2012AAFTU | 2012 | Agua Fria | 430 | 20.46 | -97.46 |
| m2012AIGRN | 2012 | Iguala | 492 | 18.35 | -99.51 |
| m2012AOBRN | 2012 | Obregon | 136 | 27.37 | -109.93 |
| m2012BAFNN | 2012 | Agua Fria | 430 | 20.46 | -97.46 |
| m2012BAL | 2012 | Almoloya | 2544 | 19.41 | -99.74 |
| m2012BBACA | 2012 | El Batan | 2281 | 19.53 | -98.85 |
| m2012BBAEF | 2012 | El Batan | 2281 | 19.53 | -98.85 |
| m2012BBANN | 2012 | El Batan | 2281 | 19.53 | -98.85 |
| m2012BCH | 2012 | G Victoria | 800 | 16.44 | -93.13 |
| m2012BCL | 2012 | Celaya | 1894 | 20.58 | -100.82 |
| m2012BEB | 2012 | Cortazar | 1770 | 20.45 | -101.04 |
| m2012BMO | 2012 | Numaran | 1735 | 20.26 | -101.93 |
| m2012BNY | 2012 | San Pedro | 1424 | 21.22 | -104.73 |
| m2012BOBRN | 2012 | Obregon | 136 | 27.37 | -109.93 |
| m2012BTORN | 2012 | Torreon | 1137 | 25.56 | -103.37 |

**Table S1** Location, year, coordinates, and elevation of each location held for phenotypic field trials.

| | ASI | BareCobWeight | DaysToFlowering | FieldWeight | GrainWeightPerHectareCorrected | PlantHeight |
|---|---|---|---|---|---|---|
| m2011BAFNN | 1279 | 1276 | 1279 | 1279 | 0 | 1278 |
| m2011BCH | 0 | 326 | 0 | 326 | 0 | 308 |
| m2011BCL | 489 | 492 | 489 | 492 | 490 | 492 |
| m2011BMO | 234 | 235 | 234 | 0 | 0 | 235 |
| m2011BNY | 402 | 400 | 402 | 0 | 0 | 401 |
| m2011BTLNN | 0 | 0 | 665 | 0 | 0 | 665 |
| m2011BTORN | 954 | 953 | 954 | 0 | 0 | 954 |
| m2012AAFCA | 1432 | 0 | 1432 | 1434 | 0 | 1434 |
| m2012AAFTU | 1434 | 0 | 1434 | 1434 | 0 | 1434 |
| m2012AIGRN | 0 | 0 | 0 | 0 | 0 | 514 |
| m2012AOBRN | 0 | 316 | 316 | 316 | 316 | 316 |
| m2012BAFNN | 424 | 0 | 424 | 0 | 0 | 424 |
| m2012BAL | 492 | 0 | 493 | 0 | 0 | 522 |
| m2012BBACA | 606 | 0 | 606 | 0 | 0 | 607 |
| m2012BBAEF | 719 | 0 | 719 | 719 | 0 | 719 |
| m2012BBANN | 607 | 606 | 607 | 0 | 0 | 607 |
| m2012BCH | 457 | 456 | 457 | 457 | 456 | 457 |
| m2012BCL | 0 | 482 | 484 | 484 | 482 | 484 |
| m2012BEB | 0 | 337 | 340 | 340 | 334 | 340 |
| m2012BMO | 0 | 144 | 144 | 0 | 0 | 144 |
| m2012BNY | 0 | 479 | 485 | 485 | 480 | 485 |
| m2012BOBRN | 0 | 440 | 431 | 0 | 0 | 440 |
| m2012BTORN | 189 | 189 | 189 | 189 | 188 | 189 |

**Table S2** Number of accessions in each trial (location and year) measured for each trait.

| | CML244/CML349 | CML269/CML264 | CML373/CML311 | CML451/CML486 | CML457/CML459 | CML495/CML494 |
|---|---|---|---|---|---|---|
| m2011BAFNN | 0 | 1964 | 370 | 1693 | 0 | 2364 |
| m2011BCH | 0 | 395 | 59 | 193 | 0 | 313 |
| m2011BCL | 0 | 696 | 1507 | 300 | 0 | 441 |
| m2011BMO | 12 | 168 | 348 | 40 | 224 | 146 |
| m2011BNY | 0 | 531 | 0 | 398 | 0 | 676 |
| m2011BTLNN | 18 | 232 | 570 | 100 | 262 | 148 |
| m2011BTORN | 0 | 947 | 1096 | 764 | 48 | 960 |
| m2012AAFCA | 0 | 1586 | 768 | 1402 | 0 | 1976 |
| m2012AAFTU | 0 | 1588 | 768 | 1404 | 0 | 1976 |
| m2012AIGRN | 0 | 123 | 87 | 84 | 0 | 220 |
| m2012AOBRN | 0 | 375 | 345 | 320 | 0 | 540 |
| m2012BAFNN | 0 | 402 | 315 | 243 | 0 | 312 |
| m2012BAL | 207 | 0 | 81 | 19 | 1200 | 0 |
| m2012BBACA | 232 | 195 | 93 | 57 | 1242 | 0 |
| m2012BBAEF | 312 | 0 | 144 | 84 | 2336 | 0 |
| m2012BBANN | 312 | 259 | 124 | 76 | 1656 | 0 |
| m2012BCH | 0 | 2254 | 0 | 168 | 0 | 318 |
| m2012BCL | 0 | 586 | 650 | 490 | 325 | 365 |
| m2012BEB | 0 | 597 | 464 | 250 | 5 | 375 |
| m2012BMO | 0 | 0 | 282 | 150 | 0 | 0 |
| m2012BNY | 0 | 583 | 650 | 489 | 322 | 370 |
| m2012BOBRN | 0 | 0 | 555 | 756 | 0 | 0 |
| m2012BTORN | 0 | 245 | 132 | 510 | 0 | 246 |

**Table S3** Number of accessions in each trial crossed to each tester, before filtering.

| | Experimento | ASI | BareCobWeight | DaysToFlowering | FieldWeight | GrainWeightPerHectareCorrected | PlantHeight |
|---|---|---|---|---|---|---|---|
| 1 | m2011BAFNN | 4 | 4 | 4 | 4 | 0 | 4 |
| 2 | m2011BCL | 4 | 4 | 4 | 4 | 4 | 4 |
| 3 | m2011BMO | 6 | 6 | 6 | 0 | 0 | 6 |
| 4 | m2011BNY | 3 | 3 | 3 | 0 | 0 | 3 |
| 5 | m2011BTORN | 5 | 5 | 5 | 0 | 0 | 5 |
| 6 | m2012AAFCA | 4 | 0 | 4 | 4 | 0 | 4 |
| 7 | m2012AAFTU | 4 | 0 | 4 | 4 | 0 | 4 |
| 8 | m2012BAFNN | 4 | 0 | 4 | 0 | 0 | 4 |
| 9 | m2012BAL | 3 | 0 | 3 | 0 | 0 | 4 |
| 10 | m2012BBACA | 5 | 0 | 5 | 0 | 0 | 5 |
| 11 | m2012BBAEF | 4 | 0 | 4 | 4 | 0 | 4 |
| 12 | m2012BBANN | 5 | 5 | 5 | 0 | 0 | 5 |
| 13 | m2012BCH | 3 | 3 | 3 | 3 | 3 | 3 |
| 14 | m2012BTORN | 4 | 4 | 4 | 4 | 4 | 4 |
| 15 | m2011BCH | 0 | 4 | 0 | 4 | 0 | 4 |
| 16 | m2012AOBRN | 0 | 4 | 4 | 4 | 4 | 4 |
| 17 | m2012BCL | 0 | 5 | 5 | 5 | 5 | 5 |
| 18 | m2012BEB | 0 | 5 | 5 | 5 | 5 | 5 |
| 19 | m2012BMO | 0 | 2 | 2 | 0 | 0 | 2 |
| 20 | m2012BNY | 0 | 5 | 5 | 5 | 5 | 5 |
| 21 | m2012BOBRN | 0 | 2 | 2 | 0 | 0 | 2 |
| 22 | m2011BTLNN | 0 | 0 | 6 | 0 | 0 | 6 |
| 23 | m2012AIGRN | 0 | 0 | 0 | 0 | 0 | 4 |

**Table S4** Number of unique testers used for crossing in each trial.

| | Tester | ASI | BareCobWeight | DaysToFlowering | FieldWeight | GrainWeightPerHectareCorrected | PlantHeight |
|---|---|---|---|---|---|---|---|
| 1 | CML244/CML349 | 5 | 2 | 6 | 1 | 0 | 6 |
| 2 | CML269/CML264 | 12 | 13 | 17 | 11 | 7 | 19 |
| 3 | CML373/CML311 | 12 | 13 | 19 | 11 | 6 | 21 |
| 4 | CML451/CML486 | 13 | 15 | 20 | 12 | 7 | 23 |
| 5 | CML457/CML459 | 6 | 6 | 10 | 4 | 3 | 10 |
| 6 | CML495/CML494 | 10 | 12 | 15 | 11 | 7 | 17 |

**Table S5** Number of trials where each tester was used for crossing, before filtering.

| | PCA1 | PCA2 | PCA3 | PCA4 | PCA5 |
|---|---|---|---|---|---|
| LatNew | 0.467347161304051 | -0.582016495540008 | -0.0263646420609473 | 0.194085817898787 | -0.100506483432986 |
| tmin | -0.695870912641895 | -0.191326687302976 | -0.137511953299172 | 0.0919695479923274 | 0.162826751154115 |
| tmax | -0.383063018300278 | -0.366988995639653 | 0.367339838652052 | -0.0705112679476859 | -0.0761503661660446 |
| trange | 0.478151335283827 | -0.0817751935172956 | 0.455532838290281 | -0.16191431249403 | -0.246984399144073 |
| precipTot | -0.401355955159194 | -0.110596096553222 | -0.116186354779725 | 0.336666133657853 | 0.171611138661601 |
| aridityMean | -0.336948056069939 | 0.0388142890979823 | -0.11218665902801 | 0.307226524904314 | 0.13841853050691 |
| rhMean | -0.308872681999844 | 0.00791112025032793 | -0.308782224887993 | 0.208268999828098 | 0.180460735501878 |
| elevation | 0.834472388150217 | 0.0235576931959703 | -0.0643642053099372 | 0.0945027976226231 | -0.0776552588685377 |

**Table S6** Correlation matrix between top 5 PCs and selected environmental variables.

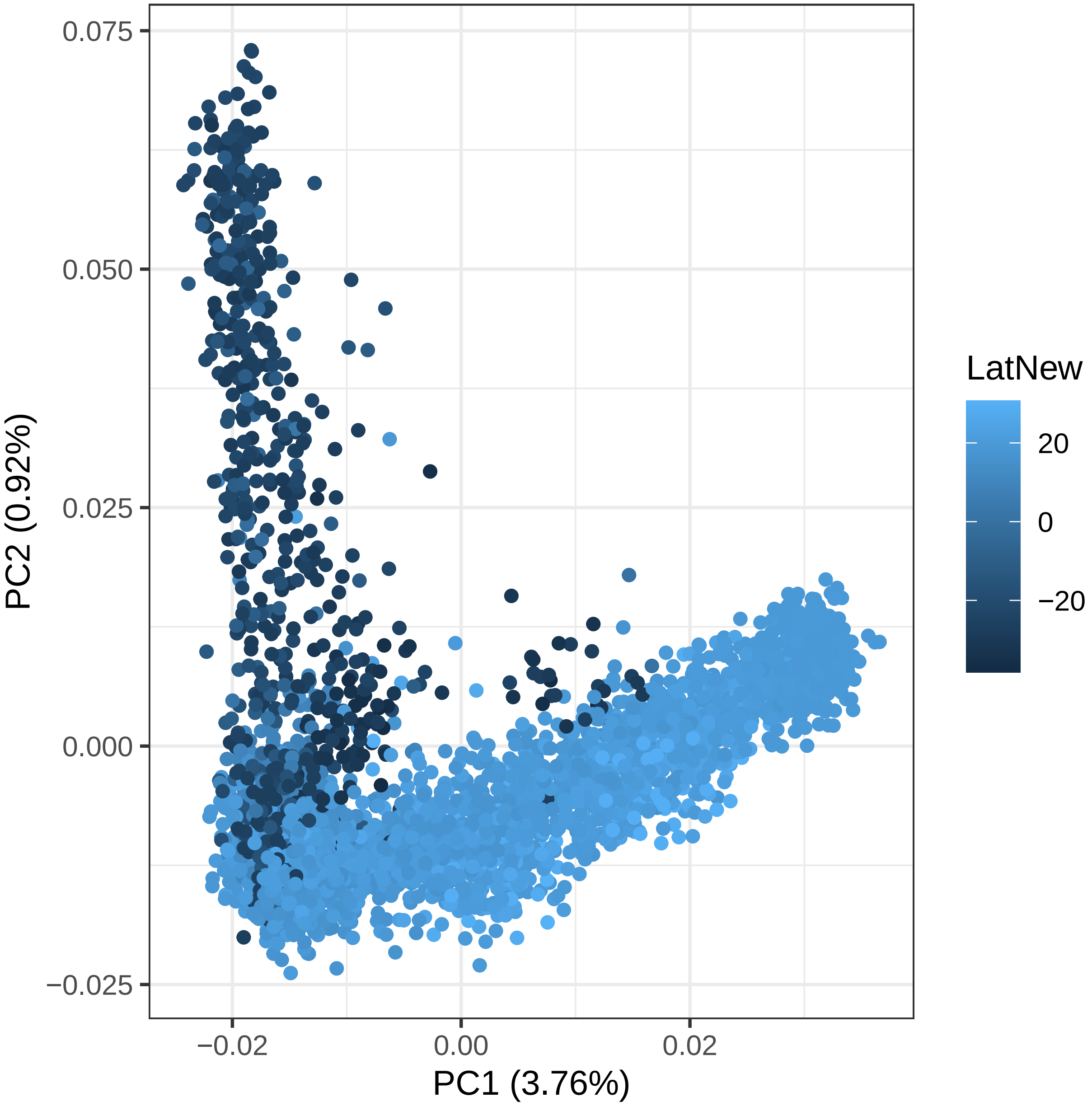


**Figure S1** Biplot of PCA1 vs PCA2 for all accessions' genotypes, colored by latitude.

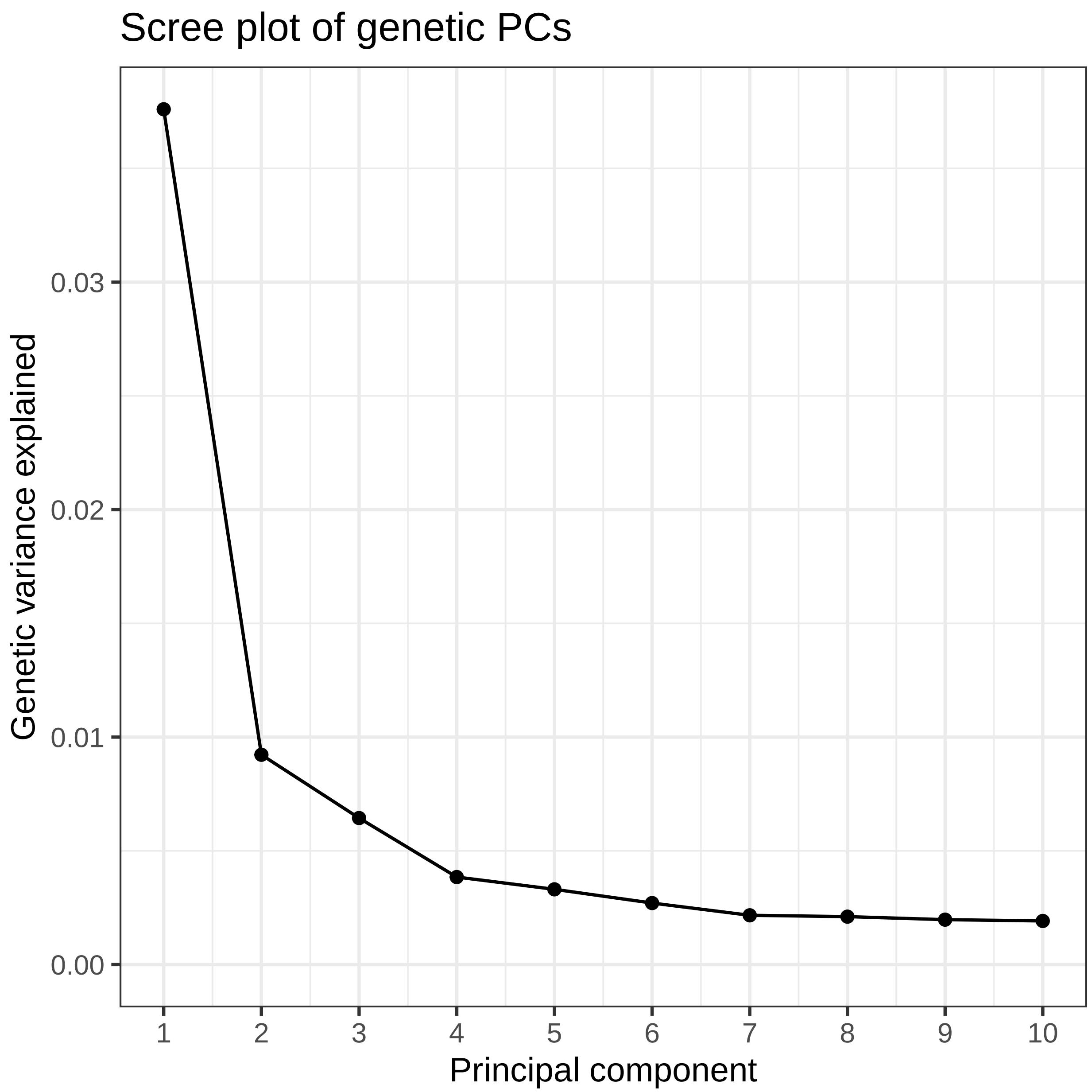


**Figure S2** Scree plot of relative contribution of top PCs towards genetic variation in SeeD population.

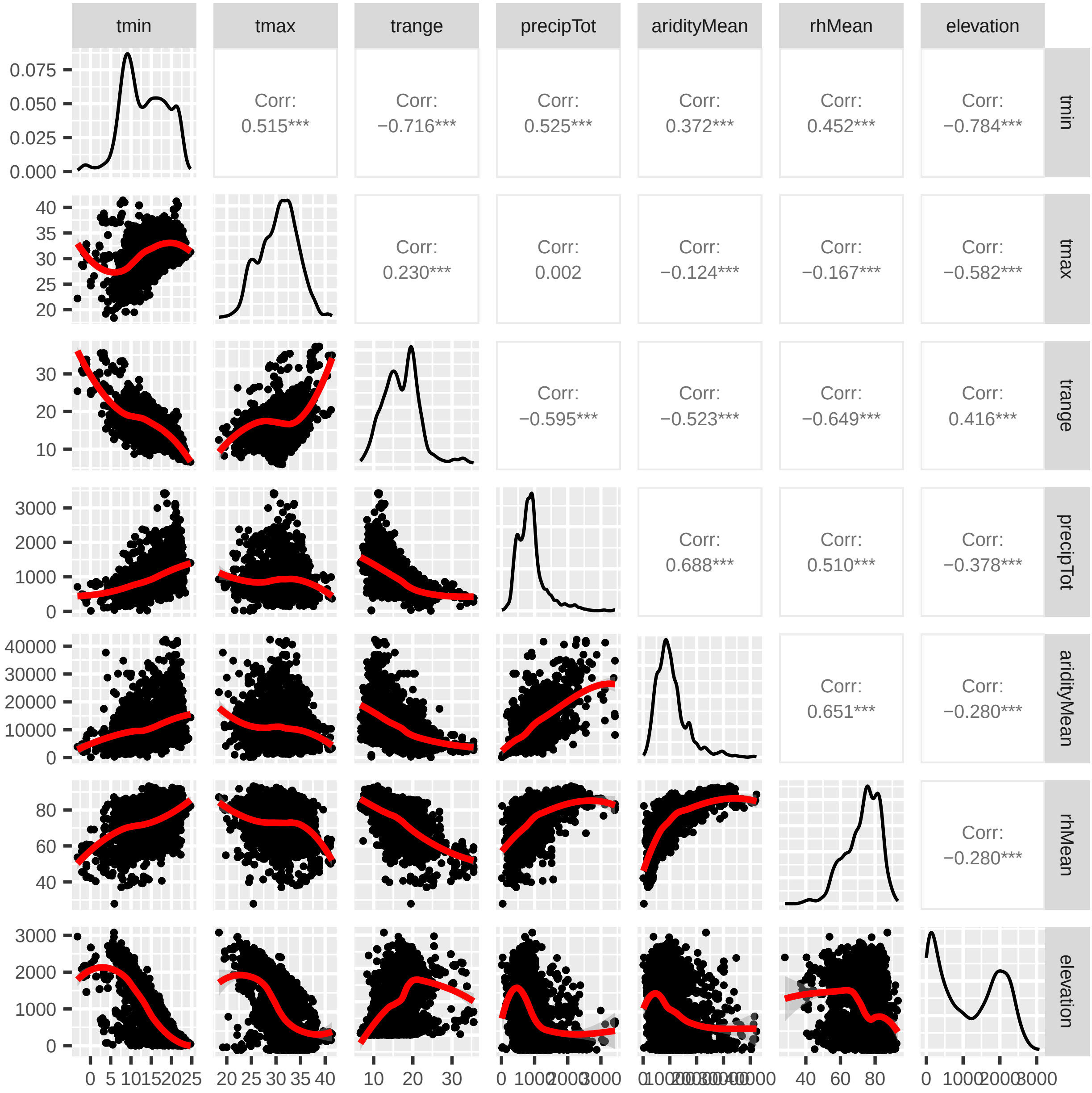


**Figure S3** Distributions and correlations between non-INT-transformed environment variables for accessions used in GEA analysis.

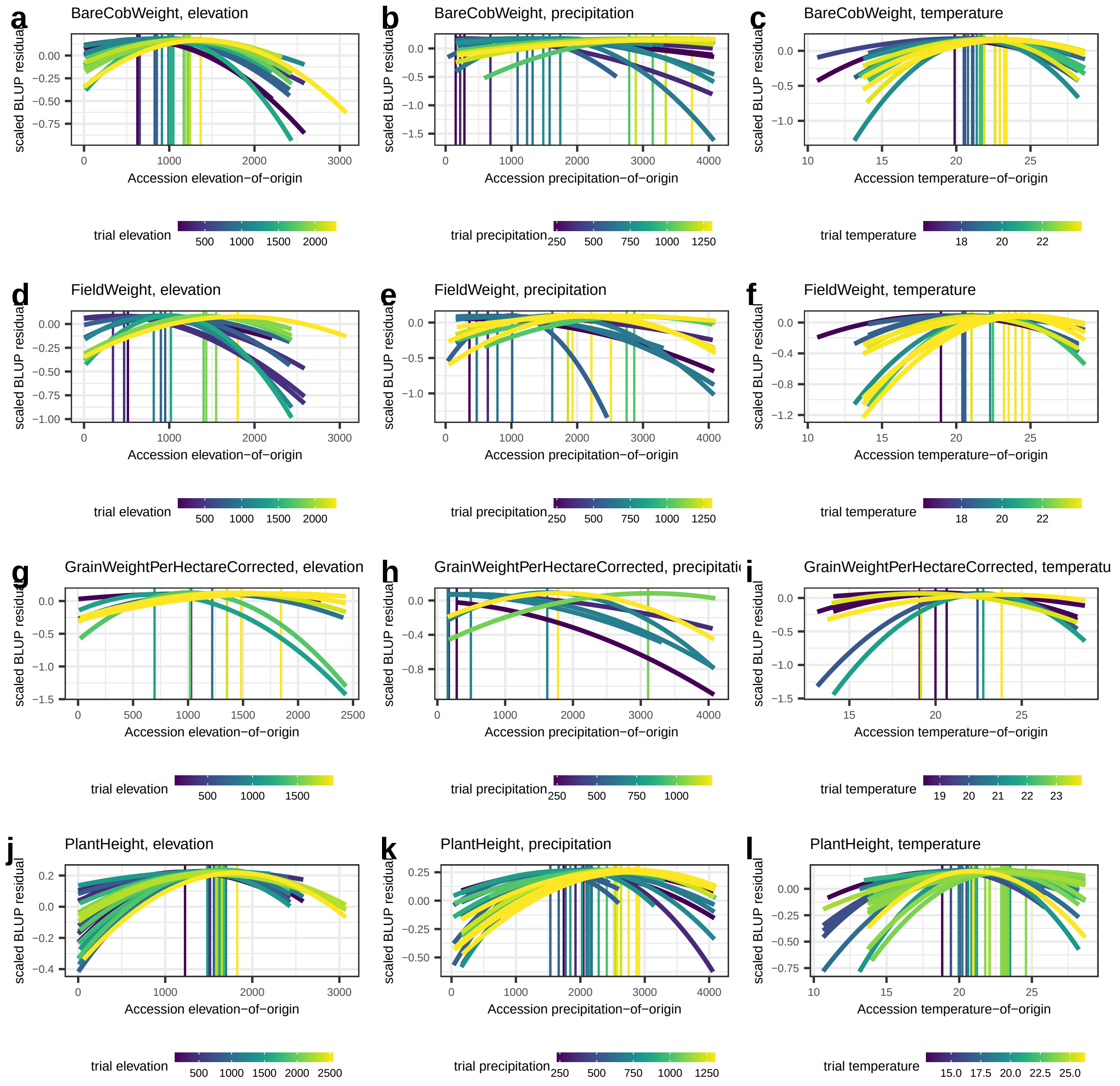


**Figure S4** All transfer plot model curves for selected yield component phenotypic trait and environmental variable combinations. Panels **a-c** measure BLUP residuals for bare cob weight, **d-f** for field weight, **g-i** for corrected grain weight, and **j-l** for plant height.

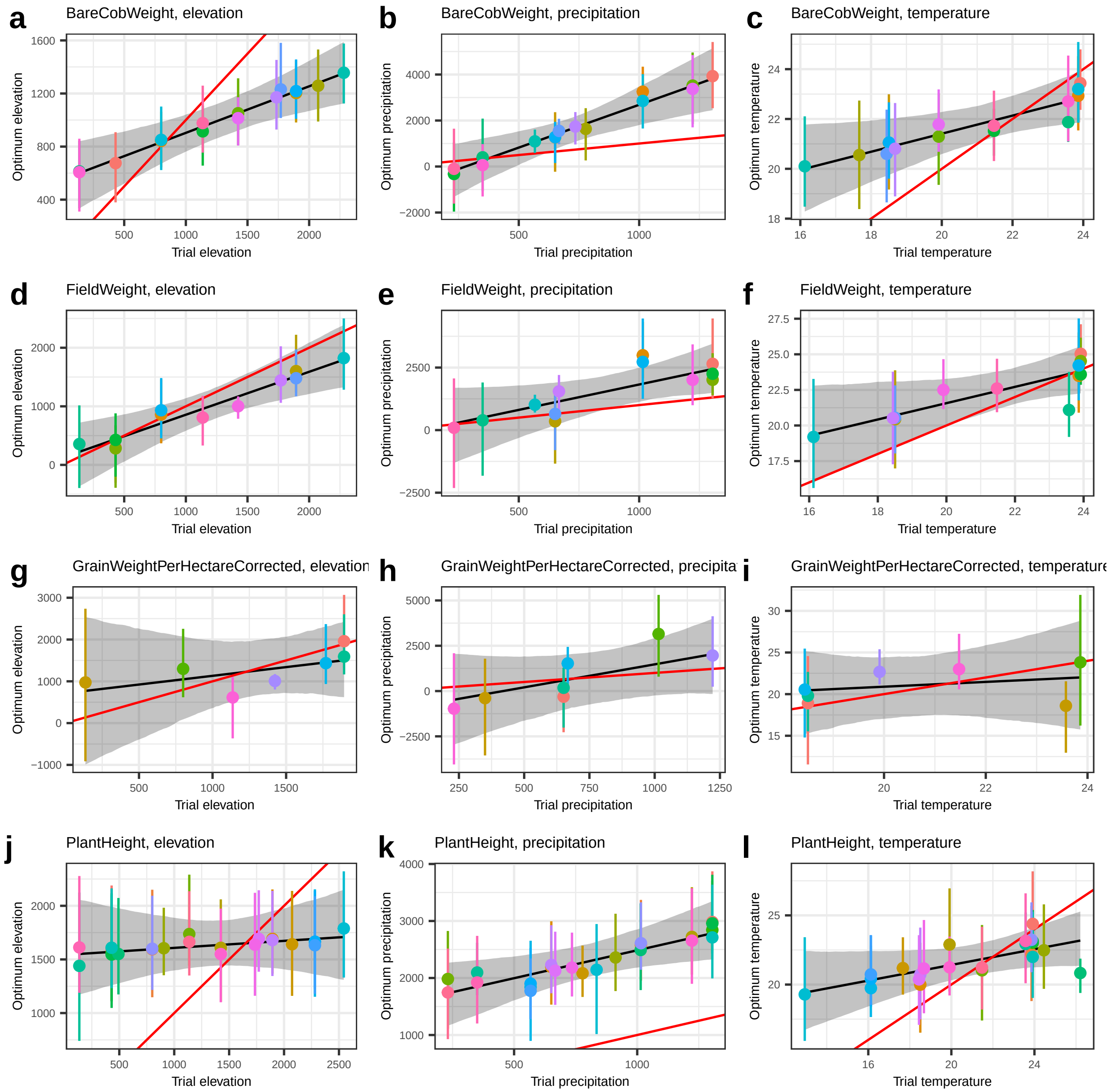


**Figure S5** All regression plots of optimal environmental value for a given trial against observed environmental value. Panels **a, d, g j** measure elevation, **b, e, h k** measure temperature, **c, f, i, l** measure precipitation. Individual dots represent trials with measurements for the specific combination.

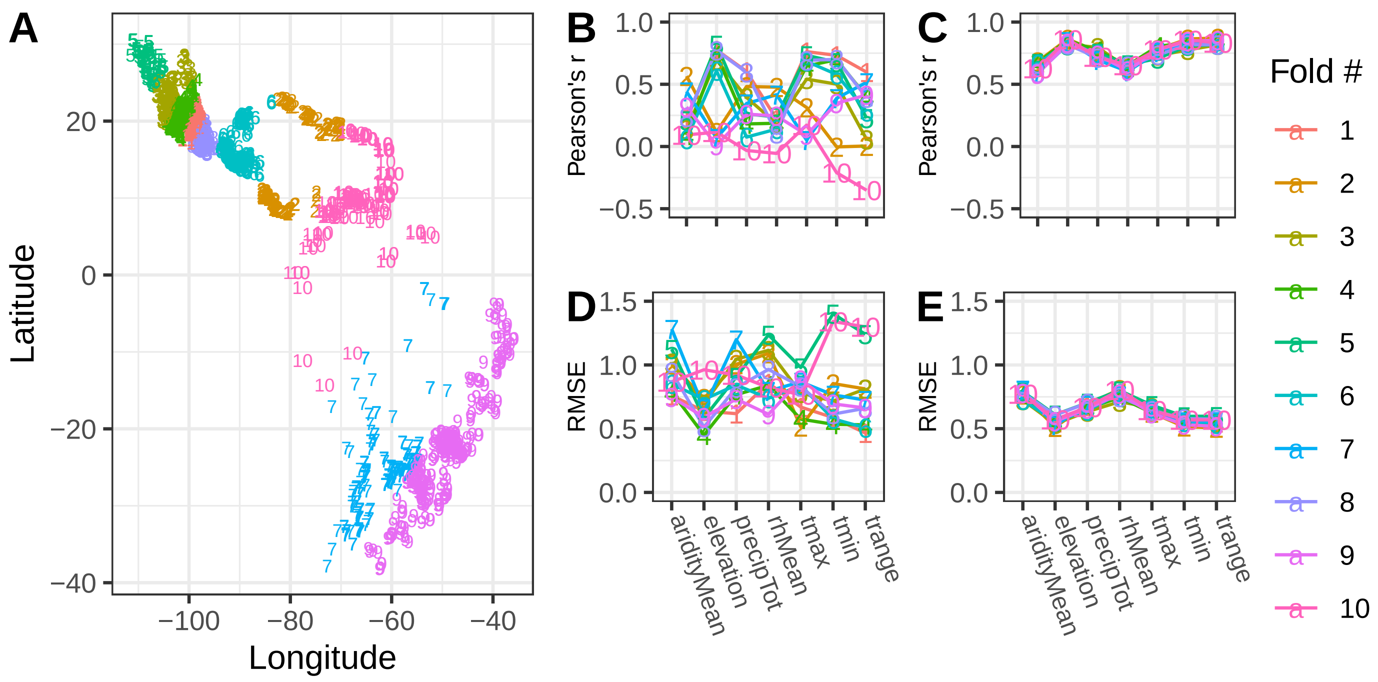


**Figure S6 Correlations between genetics and environment are demonstrated through prediction but decrease when considering spatial distance. A)** Map of sampled CV folds used in spatial GPoE. **(B-E)** Pearson's *r* correlation and RMSE for spatial (B, D) and random (C, E) sampling for each environmental variable across folds.

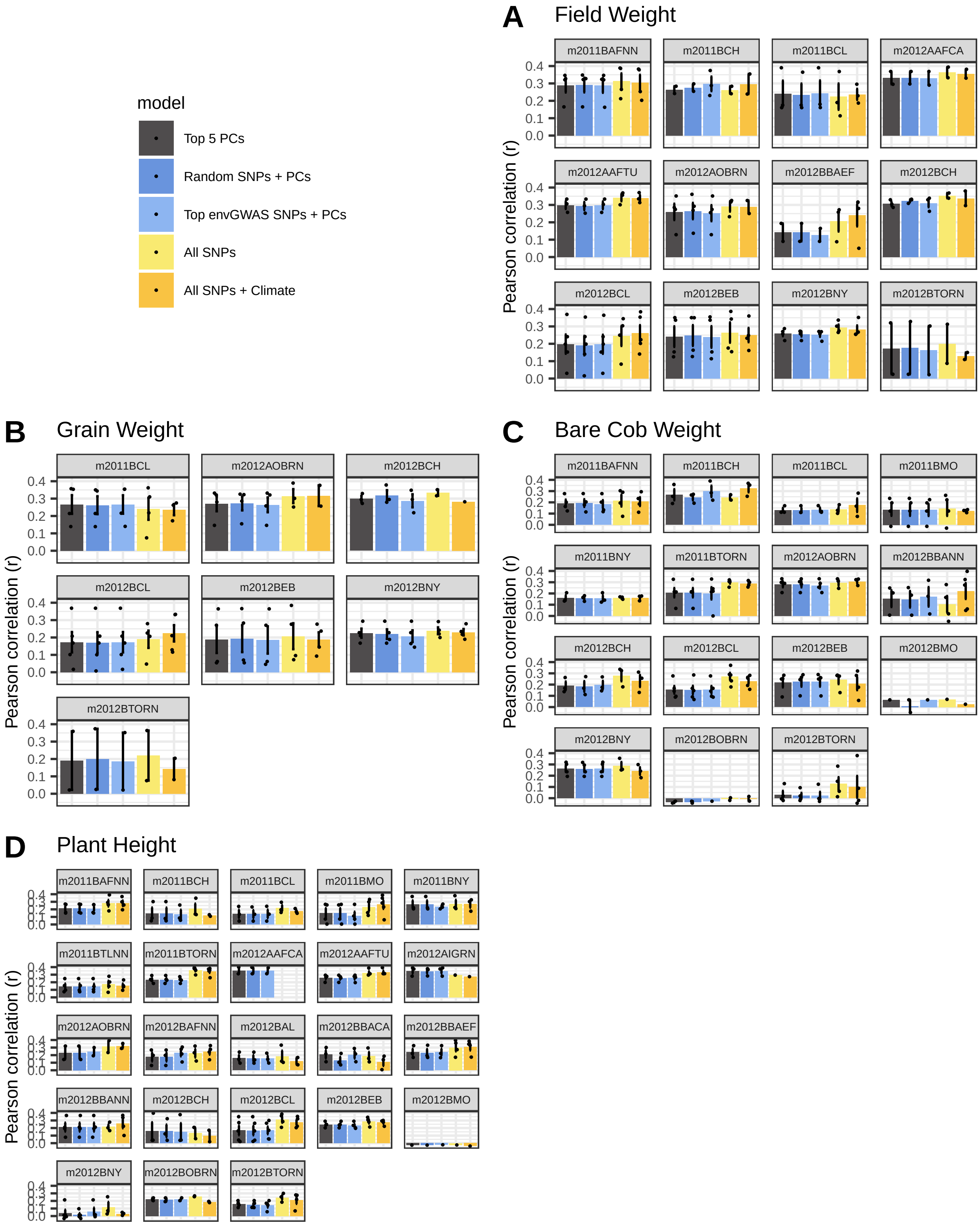


**Figure S7** Predictive ability results for field weight, corrected grain weight per hectare, bare cob weight, and plant height, where panels are separated by trial (location and year). Predictive ability for each model was measured as Pearson's r correlation to observed BLUP value. Predictions were made separately for each set of accessions crossed to a given tester for each trial (individual points). Models tested here included top five PCs, random SNPs + PCs, envGWAS SNPs + PCs, all SNPs, and all SNPs + climate data in a linear model.

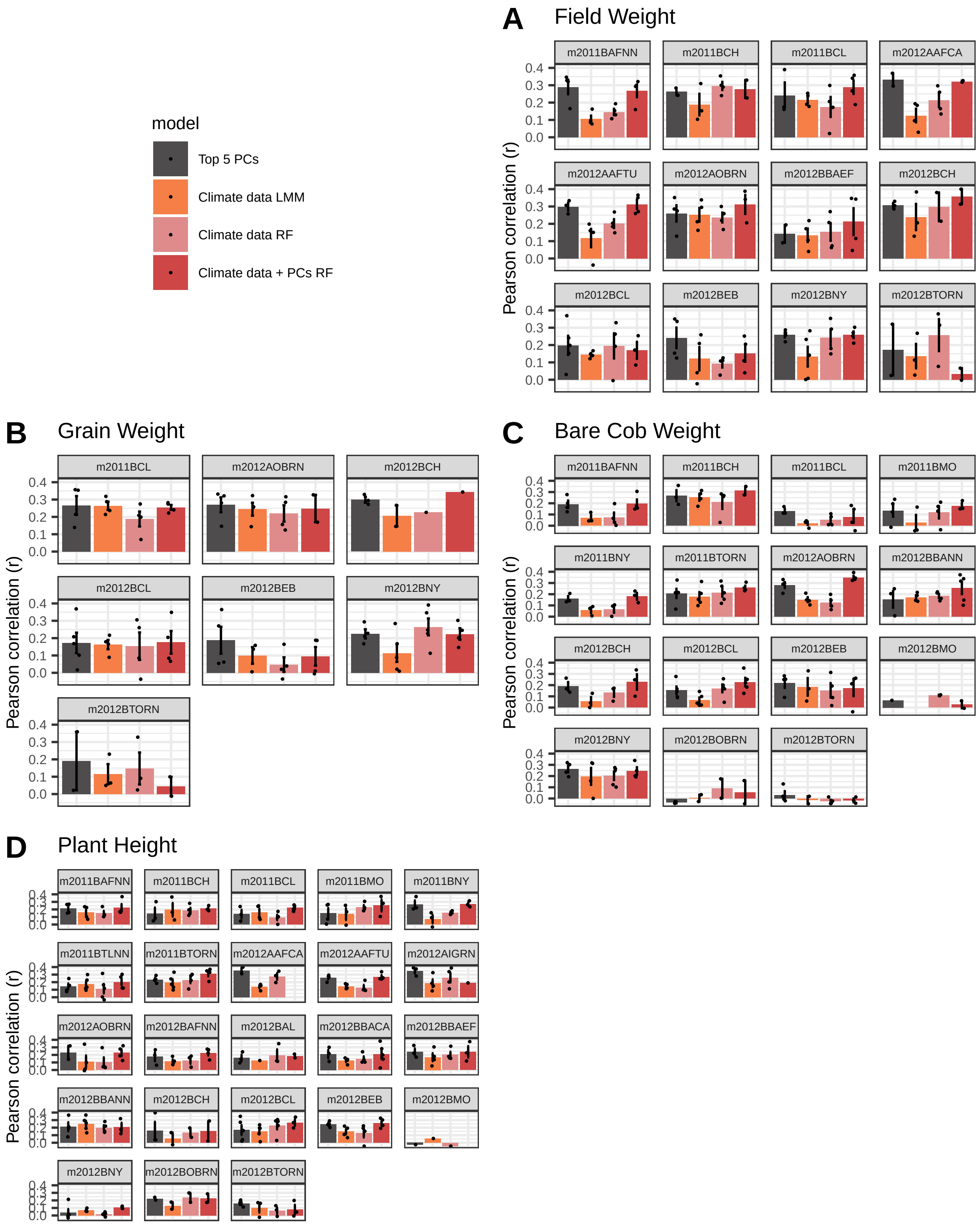


**Figure S8** Environmental data model predictive ability results for field weight, corrected grain weight per hectare, bare cob weight, and plant height, where panels are separated by trial (location and year). Predictions were made separately for each set of accessions crossed to a given tester for each trial (individual points). Models tested here included top five PCs, climate data via linear models, climate data modeled with random forests, and climate data + top five PCs with random forests.

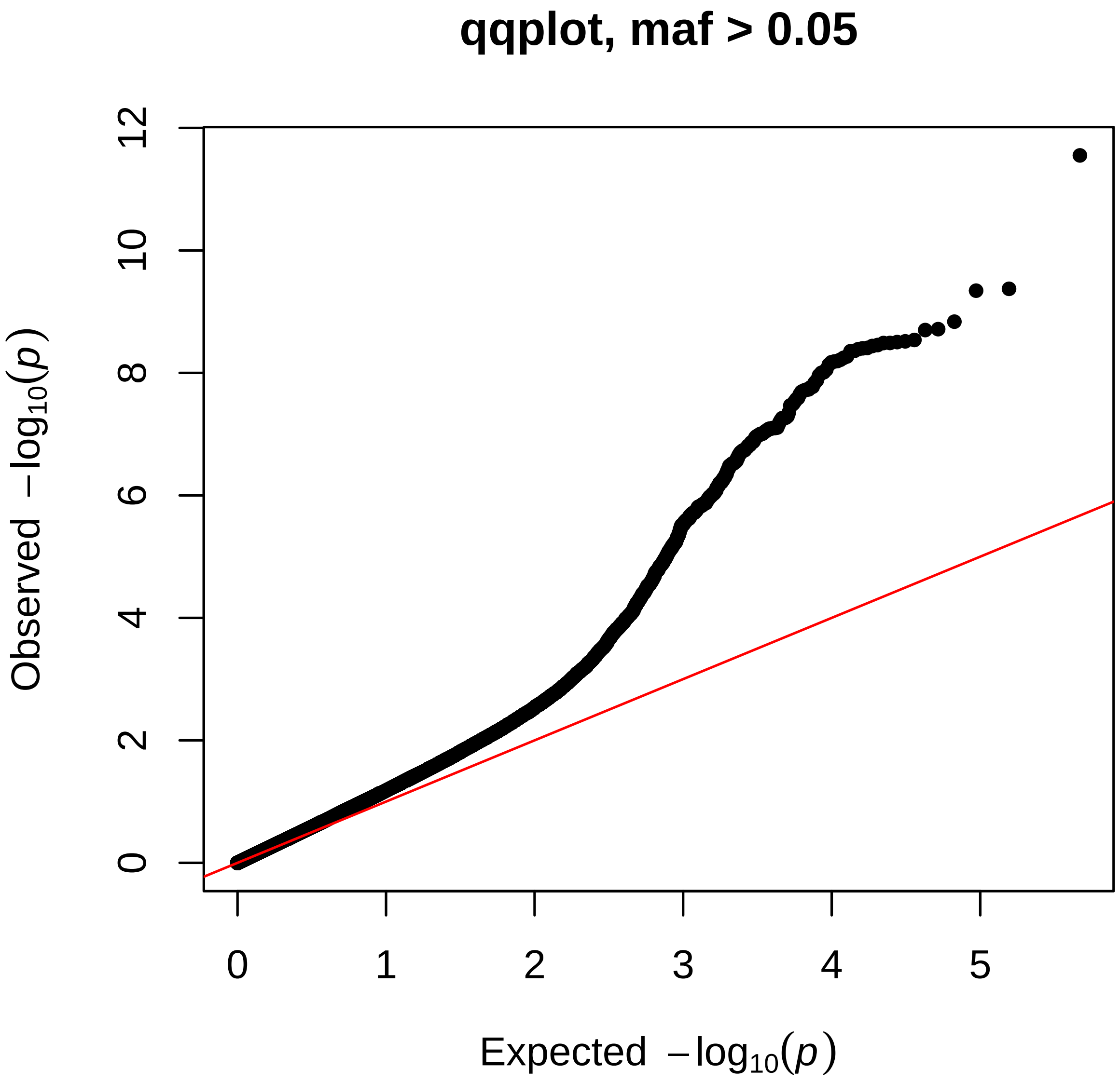


**Figure S9** QQ-plot of multivariate joint envGWAS.

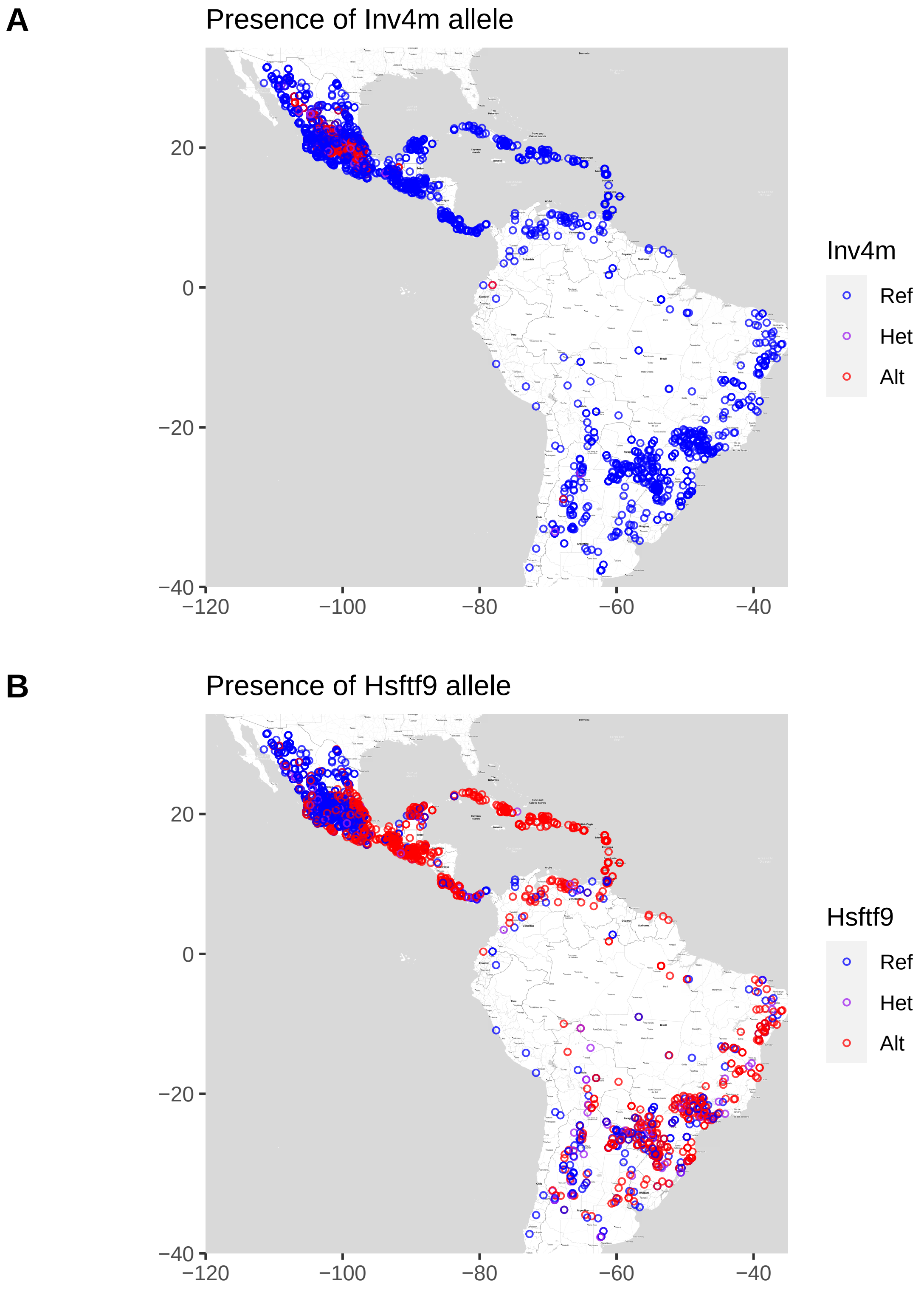


**Figure S10** Spatial distributions of lead SNPs representing **(A)** the *Inv4m* inversion and the **(B)** *hsftf9* putative candidate locus. Colors represent number of alternate allele present for given collection accession.

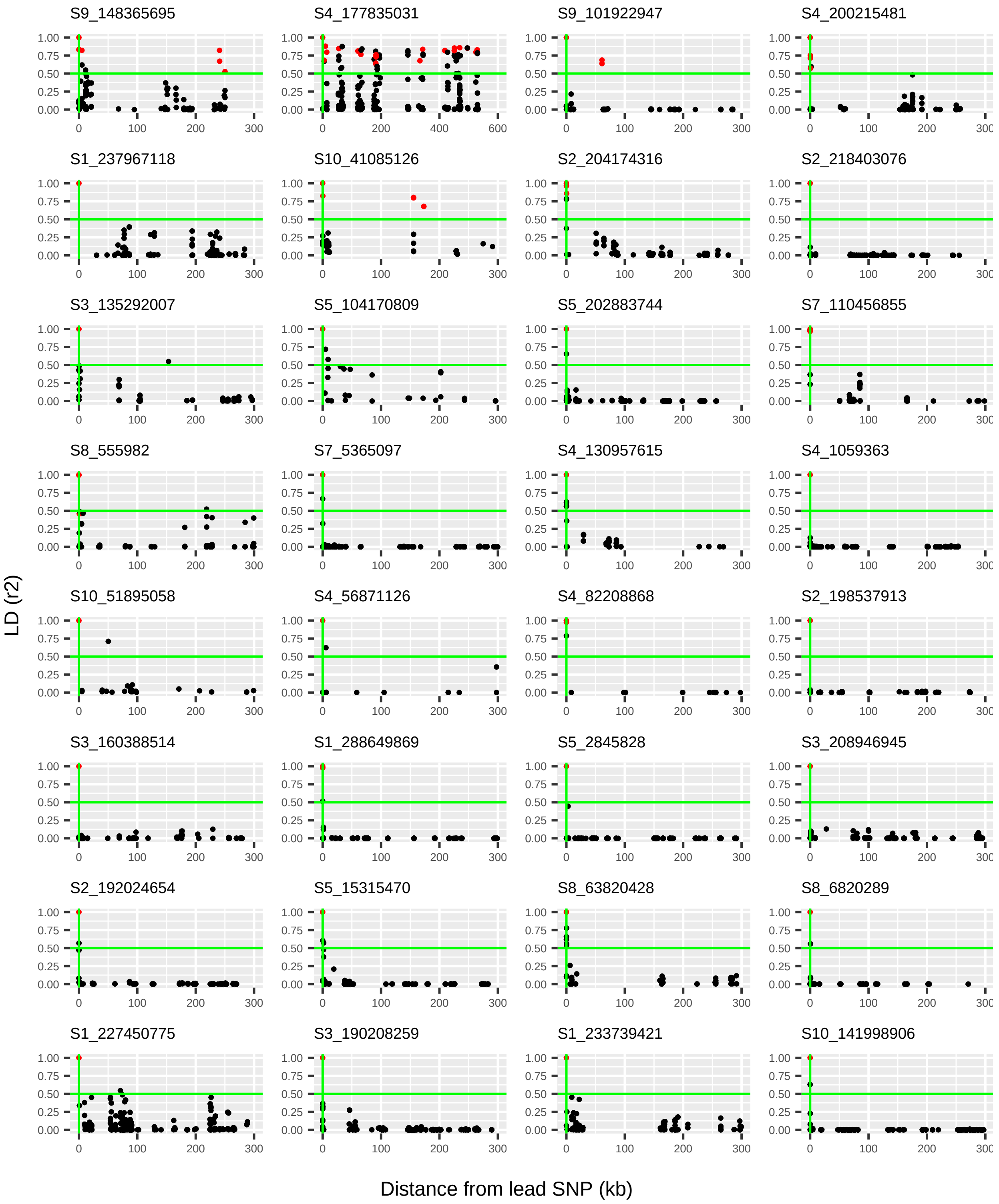


**Figure S11** Comparison of LD for all SNPs within ±300kb of top 32 lead SNPs. SNPs found significant in envGWAS are colored in red. For S4_177835031, 600kb was used to describe *Inv4m*
